# Model balancing: in search of consistent metabolic states and in-vivo kinetic constants

**DOI:** 10.1101/2019.12.23.887166

**Authors:** Wolfram Liebermeister, Elad Noor

## Abstract

Enzyme kinetic constants *in vivo* are largely unknown, which limits the construction of large metabolic models. While model fitting, in principle, aims at fitting kinetic constants to measured metabolic fluxes, metabolite concentrations, and enzyme concentrations, the resulting estimation problems are typically non-convex and hard to solve, especially if models are large. Here we assume that metabolic fluxes are known and show how consistent kinetic constants, metabolite concentrations, and enzyme concentrations can be determined simultaneously from data. If one specific term is omitted – a term that penalises small enzyme concentrations – we obtain a convex optimality problem with a unique local optimum. The estimation method with or without this term, called model balancing, applies to models with a wide range of rate laws and accounts for thermodynamic constraints on kinetic constants and metabolite concentrations through thermodynamic forces. It can be used to estimate *in-vivo* kinetic constants from omics data, to complete and adjust available data, or to construct plausible metabolic states with a predefined flux distribution. As a demonstrative case, we balance a model of *E. coli* central metabolism with artificial or experimental data. The tests show what information about kinetic constants can be obtained from omics data, and reveal the practical limits of estimating *in-vivo* kinetic constants.

## 1 Introduction

As the number of metabolic network reconstructions is constantly growing, so is the desire to convert metabolic networks into kinetic models by integration of omics data [1, 2, 3, 4, 5]. To build models with plausible parameters and metabolic states (characterised by enzyme concentrations, metabolite concentrations, and fluxes), one needs to reconstruct the metabolic network, add allosteric regulation arrows, choose the enzymatic rate laws, determine the kinetic constants, and construct plausible metabolic states. All these subproblems have been addressed in one way or another. Pathway models have been built from *in-vitro* enzyme kinetics [6, 7]. To simplify model construction and to deal with unknown rate laws, modellers proposed standardised rate laws [1, 8], used them to generate models automatically [9], and evaluated them for their practical use [10]. *In-vitro* kinetic constants, available from the Brenda database [11, 12], are widely used, and unknown *k*_cat_ values have been estimated by machine learning [13]. Inserting measured or sampled kinetic constants directly into models can lead to inconsistencies, given that thermodynamics implies dependencies between kinetic constants via Wegscheider conditions and Haldane relationships. To address this problem, methods to construct thermodynamically consistent parameter sets have been devised [14, 15, 16, 17, 18] and applied in modelling [5]. In parallel, there have been attempts to estimate *in-vivo* kinetic constants [19] from flux, metabolite, and enzyme data [20, 21]. Methods for parameter fitting have been developed and benchmarked [22, 23], and the question of parameter identifiability has been addressed [24]. Large models have been parameterised [2, 25], and workflows for model parameterization have been developed [26, 27]. Finally, even if parameters are unknown, parameter sampling and ensemble modelling [28, 29, 30, 31, 32] allow to find plausible parameter sets [33] and to draw conclusions about dynamic behaviour independently of specific parameter values [34].

Here we address the problem of finding realistic, consistent values of kinetic constants. The values *in vivo* are hard to measure, and the *in-vitro* values can be unreliable. To obtain *in vivo* kinetic constants, we consider a kinetic metabolic model that contains these constants as model paramaters and fit the model to measured omics data. In an ideal case in which enzyme concentrations, metabolite concentrations and fluxes are precisely known for a sufficient number of metabolic states, we can directly solve for the kinetic constants [21]. Likewise, in an ideal case in which kinetic constants and enzyme concentrations are already known precisely, metabolite concentrations and fluxes could be computed by simulating the model. In reality, we are in between these two cases: data of different types may be available, but all of them are incomplete and noisy, and cannot be used directly in models. For a complete picture and reliable estimates, these heterogeneous data must be combined – but how? In practice, this depends on our aims: in model construction we may have different aims, for example (i) finding consistent kinetic constants in plausible ranges; (ii) adjusting and completing a data set of measured kinetic constants to obtain consistent model parameters; (iii) estimating *in-vivo* kinetic constants from omics data (measured enzyme concentrations, metabolite concentrations, and fluxes); (iv) completing and adjusting omics data for consistent metabolic states, which may involve predictions of physilogical metabolite concentrations [35, 36] or predictions of metabolite and enzyme concentrations based on resource allocation principles [37, 38, 39].

Most generally, our task is not only to fit kinetic constants, but also to reconstruct consistent metabolite concentrations, enzyme concentration, and metabolic fluxes, based on available data for all these quantities. Since the data will be uncertain, inconsistent and incomplete, the uncertainties in data and estimated state variables and kinetic constants need to be quantified. Luckily, we can reduce these uncertainties by imposing further constraints by physical laws, including thermodynamic relations (Wegscheider conditions and Haldane relationships) between kinetic constants [1, 8] and relations between kinetic constants and metabolic variables (e.g. the fact that flux directions follow from equilibrium constants and metabolite concentrations). In addition, we can use prior distributions (for kinetic constants, metabolite concentrations, or enzyme concentrations); and we can use measured (*in vitro*) parameter values as additional data (of course, these data cannot be used as test data anymore). However, one problem remains. The resulting estimation problems typically lead to non-convex optimality problems with multiple local optima: global optima may be hard to find especially if networks are large.

Here we address, as a general problem, the estimation of consistent kinetic constants and metabolic states based on heterogenous (kinetic, metabolomics, and proteomics) data, assuming that the metabolic fluxes are given (from measurements or previous calculations) and thermodynamically consistent. We show that parameter fitting in kinetic metabolic models can be formulated, with the exception of one term that penalises small enzyme concentrations, as a convex optimality problem. The resulting estimation method, called model balancing, uses the following input data: measured or assumed values of kinetic constants (which may be incomplete and uncertain), measured or assumed values of metabolite and enzyme concentrations in a number of metabolic states (which may be incomplete and uncertain), and metabolic fluxes from the same metabolic states (which may be stationary as in FBA, or non-stationary as in dynamic simulation of a kinetic model^1^). From these data it determines a set of kinetic parameters and state variables (metabolite and enzyme concentrations for all metabolic states in question) that are consistent with the rate laws and all other dependencies in the model, plausible (i.e. respecting constraints and prior distributions), and resemble the data (showing large likelihood values). The only mandatory data are the metabolic fluxes (while other data are optional, but can help make the estimates more precise). For the estimation, we maximise the Bayesian posterior density (maximum-likelihood estimation can be obtained as a special case by the use of flat priors). If we neglect one specific term (a part of the posterior that penalises small enzyme concentrations) and assume non-flat priors, the maximum-posterior problem has a single solution which can be found be gradient descent. This convexity relies on two main assumptions: (i) all fluxes are predefined and thermodynamically consistent (i.e. infeasible cycle fluxes must be excluded) and (ii) kinetic constants and metabolite concentrations are described in log-scale. Model balancing builds upon some previous methods for model construction: parameter balancing [15, 18], elasticity sampling [40], and enzyme cost minimisation [39], which we review in the discussion section.

## 2 Parameter estimation in kinetic models

A programme for finding kinetic constants and metabolic states in kinetic models by Bayesian estimation was outlined in [14]. Here we follow this programme and show that it becomes mathematically more convenient if fluxes are given (using insights about convexity from [39]).

### 2.1 Estimating kinetic constants from omics data

How can information about *in-vivo* kinetic constants be obtained from omics data? Let us first have a look at an existing approach. To estimate *k*_cat_ values, Davidi *et al*. [20] compared flux data obtained from flux balance analysis (FBA) to measured proteomics, without any knowledge of metabolite concentrations or kinetic laws. The method, which was later called kinetic profiling [41], assumes (unknown) rate laws of the general form *v* = *e k*(**c**) (with flux *v* and enzyme concentration *e*), where the catalytic rate *k* depends on (unknown) metabolite concentrations (vector **c**) and varies between zero and a maximal value *k*_cat_ (called turnover rate or catalytic constant). To determine *k*_cat_ from data, we consider a cell in different metabolic states and compute the empirical catalytic rates 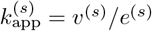 in all of these states. We can know for sure that 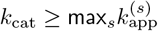. If we further assume that each enzyme reaches its maximal capacity in at least one of these states, the *k*_cat_ value (or at least, a lower bound on the *k*_cat_ value) is estimated by taking their maximum value. The method was applied to a large number of enzymes in *E. coli* and a certain correlation between the estimated *k*_cat_ values and measured *in-vivo* values was found. A part of the remaining deviations could be explained by enzyme kinetics and thermodynamics, but this was not quantitatively modelled. The limitations of this method are clear: since the max function is only sensitive to the highest value, a single high outlier value can completely distort the result. Such outliers may arise if a small protein concentration, due to measurement errors, appears even smaller. But aside from this practical problem, we also cannot be sure that enzymes reach (or come close to) their maximal capacity in one of the samples. If the estimated *in-vivo k*_cat_ value is seen as a lower bound *k*_cat_ ≥ max_*s*_ *v*^(*s*)^*/e*^(*s*)^, then how far is this bound from the true *in-vivo k*_cat_ value?

If we manage to model the (non-maximal) catalytic rates in the different states as a kinetic effect, can we maybe obtain a better estimate of the *k*_cat_ value, even if this value is not reached in any of the samples? To do this, we need to consider metabolite concentrations and enzyme kinetics, i.e. a functional form of *k*(**c**). A typical form of for a uni-uni reaction, the Michaelis-Menten kinetics, reads 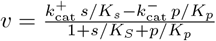 [8] or in factorised form [42] 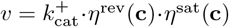 The efficiency terms *η*^rev^ (for reversibility, or thermodynamics) and *η*^sat^ (for enzyme saturation and allosteric regulation) are numbers between 0 and 1 depending on metabolite concentrations. If the efficiency terms are close to 1, then *k*(**c**) approaches its maximal value *k*_cat_; but normally *k*(**c**) is lower. Given the rate laws, and given data for fluxes **v**, metabolite concentrations **c**, and enzyme concentrations **e**, we might be able to estimate *k*_cat_ and *K*_M_ values, even if the maximal efficiency is not reached in any of the samples.

### 2.2 Metabolic model and statistical estimation model

Let us now state the estimation problem (see Figure 1 (a), for a list of mathematical symbols see table 1). We consider a kinetic metabolic model with thermodynamically consistent modular rate laws [8] and kinetic constants (e.g. equilibrium constants^2^, catalytic constants and Michaelis-Menten constants) in a vector **q** = ln **p**. We further assume a number of metabolic states, each characterised by a flux vector **v**^(*s*)^, a metabolite concentration vector **c**^(*s*)^, and an enzyme concentration vector **e**^(*s*)^. These states can be stationary (with steady-state fluxes) or non-stationary (e.g. snapshots from a dynamic time course). The model defines dependencies among model parameters and state variables, and also the kinetic constants themselves are interdependent because of physical laws [1, 8]: each rate law contains a forward and a backward catalytic constant as well as Michaelis-Menten constants, and all these parameters may depend on each other – within and across reactions – via Haldane relationships and Wegscheider conditions, which necessarily creates correlations between these constants. Activation and inhibition constants, which may also appear in the rate laws, are independent of all other kinetic constants.

**Figure 1.**
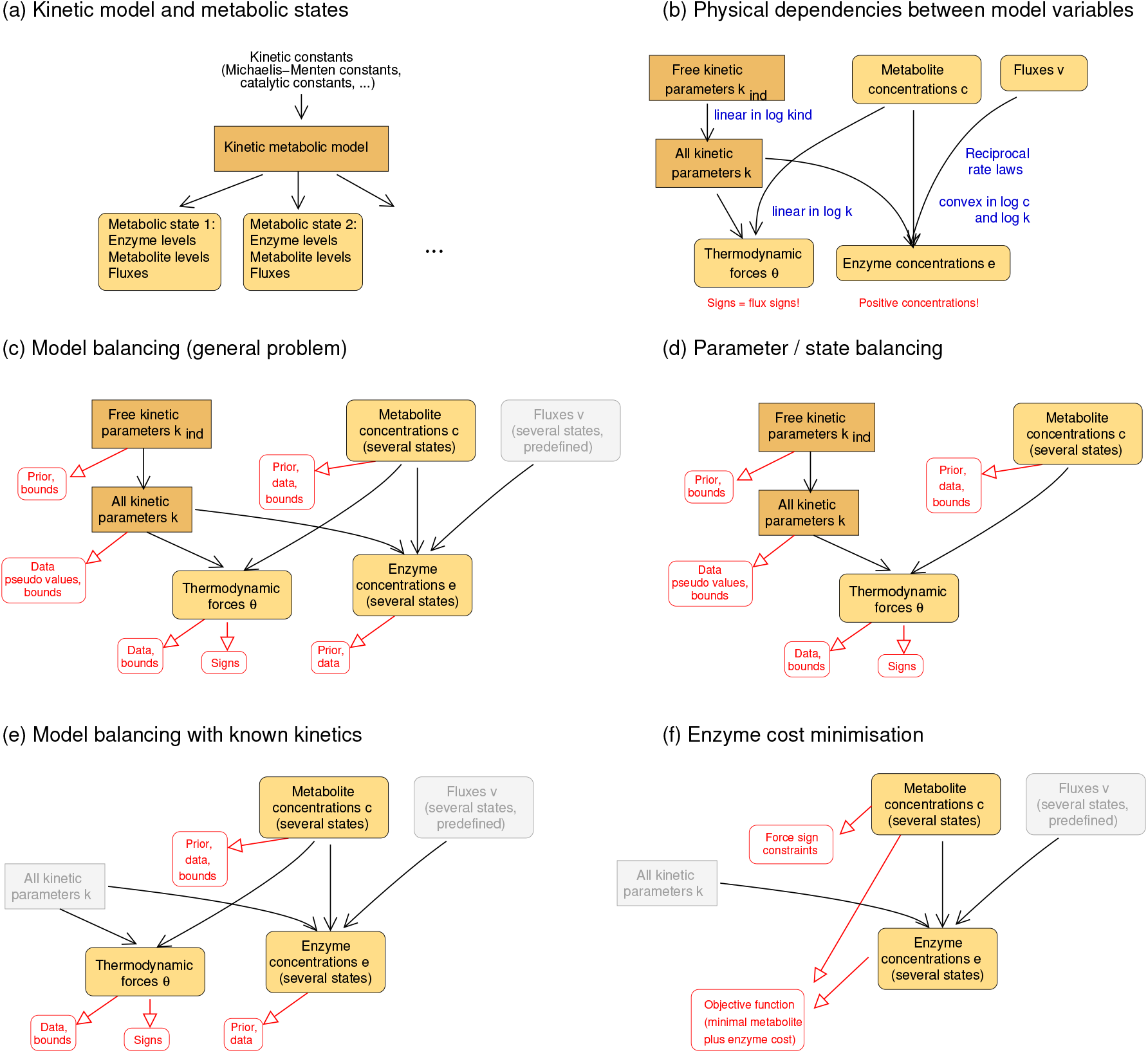
Estimation of model variables in kinetic metabolic models. (a) Kinetic model and metabolic states. A model is parameterised by kinetic constants (e.g. equilibrium constants, catalytic constants, and Michaelis-Menten constants) and gives rise to a number of metabolic states (characterised by enzyme concentrations, metabolite concentrations, and fluxes). These states may be stationary (with steady-state fluxes) or not (e.g. states during dynamic time courses). (b) Dependencies between kinetic constants and state variables. All kinetic constants are described in logarithmic scale. A subset of kinetic constants determines all other kinetic constants through linear relationships. If kinetic constants, metabolite concentrations, and fluxes are known, the enzyme concentrations can be computed from rate laws and fluxes: each enzyme concentration is a convex function of the (logarithmic) kinetic constants and metabolite concentrations. The signs of thermodynamic forces are constrained by the flux directions. (c) Parameter estimation. Kinetic constants and metabolite concentrations (for a number of metabolic states) serve as free variables of a statistical model. Dependent kinetic constants, thermodynamic driving forces, and enzyme concentrations (bottom) are treated as dependent variables, and the fluxes (top right) are predefined. For estimating the variables, priors and available data may be used. The other graphics show estimation and optimisation methods for other scenarios, in which (d) only kinetic data are balanced (no state data), (e) only state data are balanced (kinetic parameters are predefined), or (f) enzyme and metabolite concentrations are optimised for a low enzyme or metabolite cost.

**Table 1:** Mathematical symbols used in model balancing

| State variables |  |  |
| --- | --- | --- |
| Metabolic flux | $v_l^{(s)}$ | Reaction $l$, in state $s$ |
| Metabolic log-concentration | $x_i^{(s)} = \ln c_i^{(s)}$ | Metabolite $i$, in state $s$ |
| Enzyme log-concentration | $z_l^{(s)} = \ln e_l^{(s)}$ | Reaction $l$, in state $s$ |
| Kinetic constants |  |  |
| Log kinetic constants | $\mathbf{q}, \mathbf{q}^{\text{ind}}, \mathbf{q}^{\text{dep}}$ | |
| Equilibrium constants | $\mathbf{k}_{\text{eq}}, \mathbf{k}_{\text{eq}}^{\text{ind}}, \mathbf{k}_{\text{eq}}^{\text{dep}}$ | |
| Catalytic constants | $k_{\text{cat},l}^+, k_{\text{cat},l}^-$ | Reaction $l$ |
| Velocity constants | $K_{V,l} = \sqrt{k_{\text{cat},l}^+, k_{\text{cat},l}^-}$ | Reaction $l$ |
| Michaelis-Menten constants | $K_{M,li}$ | Reaction $l$, reactant $i$ |
| Activation constants | $K_{A,li}$ | Reaction $l$, activator $i$ |
| Inhibition constants | $K_{I,li}$ | Reaction $l$, inhibitor $i$ |
| Estimation score functions |  |  |
| Prior prescore | $P^*(\mathbf{x}, \mathbf{z}, \mathbf{q})$ | |
| Likelihood prescore | $L^*(\mathbf{x}, \mathbf{z}, \mathbf{q})$ | |
| Posterior prescore | $R^*(\mathbf{x}, \mathbf{z}, \mathbf{q})$ | |
| Prior score | $P(\mathbf{x}, \mathbf{q}^{\text{ind}})$ | |
| Likelihood score | $L(\mathbf{x}, \mathbf{q}^{\text{ind}})$ | |
| Posterior score | $R(\mathbf{x}, \mathbf{q}^{\text{ind}})$ | |

The dependencies between model variables are summarised in appendix A.1. To satisfy all parameter dependencies, we introduce a set of independent kinetic parameters (independent equilibrium constants, Michaelis-Menten constants, and velocity constants) that determine all remaining kinetic constants (Figure 1 (b), top left). The vector **p** contains all kinetic constants. In each metabolic state *s*, the rate laws determine the fluxes 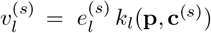 as functions of enzyme concentrations 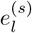, metabolite concentrations 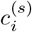, and catalytic rates *k*_*l*_. By inverting this equation, the enzyme concentrations can be written as functions

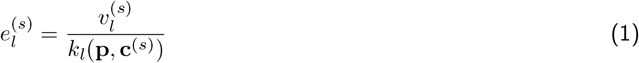

of kinetic constants, metabolite concentrations, and fluxes (Figure 1 (b), bottom). Importantly, these functions, as well as their logarithms, are convex in the space spanned by ln **p** and ln **c**^(*s*)^. The thermodynamic forces, in the vector ***θ***^(*s*)^ = ln **k**_eq_ − **<u>N</u>**^T^ ln **c**^(*s*)^, determine the flux directions (but note that vanishing fluxes are always allowed):

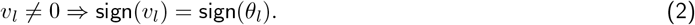

This law holds for all thermodynamically feasible rate laws.

Given the kinetic constants, state variables, and dependencies in our model, we can now define estimation problems. Our most general aim is to estimate kinetic constants, metabolite profiles and enzyme profiles in a number of metabolic states, based on data for kinetic constants, fluxes, metabolite and enzyme concentrations (and possibly thermodynamic forces) in these states. All these variables^3^, together with their dependecnies, can be visualised in a dependency schema as in Figure 1. All data may be uncertain and incomplete, except for the fluxes, which are assumed to be known. Besides dependencies and data, we may use prior distributions and impose upper and lower bounds on the model parameters and on metabolite and enzyme concentrations. In model balancing, we treat independent kinetic constants and metabolite concentrations as free variables, while and dependent kinetic constants, enzyme concentrations and thermodynamic forces are dependent variables (computed from kinetic constants, metabolite concentrations, and fluxes). All kinetic constants and concentrations – but not the thermodynamic forces and fluxes – are treated in log-scale, and natural logarithms are used throughout. The vector of free variables (kinetic constants and metabolite concentrations) is constrained by Eq. (2) and by lower and upper bounds, and the resulting solution space is a convex polytope. For a Bayesian parameter estimation, here we determine the posterior mode, but the formalism can also be readily used for maximum-likelihood estimation and posterior sampling [43]. Formulae for these problems are summarised in appendix A.1.

### 2.3 State balancing: fitting of metabolite and enzyme concentrations

Before we come to the full model balancing problem, let us first consider a simplified problem in which the kinetic constants are known and the aim is to find consistent metabolite and enzyme concentrations for a single steady state with known fluxes^4^, and given incomplete, uncertain data for some or all metabolite and enzyme concentrations. To fit consistent metabolite and enzyme concentrations, we maximise their posterior density. Metabolite and enzyme concentrations^5^ are described by their natural logarithms^6^. For the metabolite log-concentration vector **x** = ln **c**, we assume Gaussian priors^7^ (with mean vector 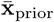 and covariance matrix 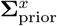) and lower and upper bounds (possibly different for each metabolite). Likewise, for the enzyme log-concentration vector **z** = ln **e**, we assume Gaussian priors (with mean vector 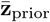 and covariance matrix 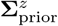). Both for metabolites and enzymes, negative concentrations are excluded^8^. The possible metabolite log-profiles **x** form a convex polytope 𝒫_**x**_ in log metabolite space [39]. The shape of this polytope is defined by physiological upper and lower bounds and by thermodynamic constraints, depending on flux directions and equilibrium constants. The metabolite log-concentrations *x*_*i*_, our free variables, determine the enzyme concentrations *e*_*l*_ through Eq. (1) and both the absolute and the enzyme log-concentrations are convex functions on the metabolite polytope (appendix section A.2). To define an estimation problem, we construct the polytope as our solution space, consider prior, likelihood and posterior functions on this polytope, and use them to estimate metabolite concentrations and corresponding enzyme concentrations.

The Bayesian estimation of model variables follows the approach outlined in [14]. Starting from the Gaussian prior density *p*(**x**) for metabolite log-concentrations **x**, we take the logarithms, omit constant terms, and obtain the *prior loss score*

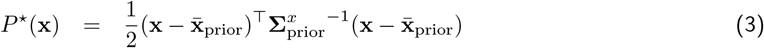

for metabolite log-concentrations. Writing quadratic functions written in short as 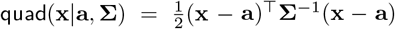, we obtain the joint prior for metabolite log-concentrations **x** = ln **c** and enzyme concentrations **z** = ln **e**

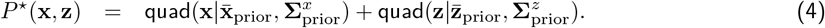

The choice of prior means and covariances is discused at the end of section 3.3. Similarly, using data for **x** and **z**, we define the *likelihood loss score*

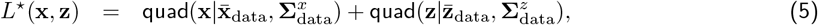

the negative log-likelihood (again without constant terms and the prefactor). The vectors 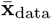, and 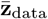 and matrices 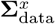 and 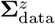 contain mean values and covariances for measurement data. For simplicity, we assume that each state variable has been measured exactly once^9^. By adding the two functions (4) and (5), we obtain the *posterior loss score R*^*\**^(**x, z**) = *P*^*\**^(**x, z**) + *L*^*\**^(**x, z**), and by simplifying the quadratic functions [14, 15], we obtain the formula

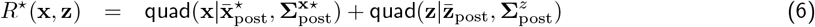

with metabolite covariance matrix and mean vector^10^

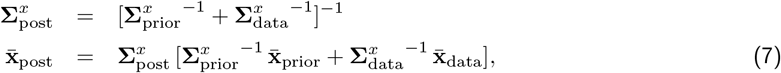

and analogous formulae for the enzyme covariance matrix 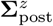 and mean vector 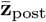. Until this point, all functions considered are convex: the prior score *P*^*^, likelihood score *L*^*\**^, and posterior score *R*^*\**^ are convex in **x** and **z**, and if non-flat priors are used, then prior and posterior score are strictly convex.

Now, as we remember, the enzyme concentrations are not free variables, but fully dependent on metabolite concentrations and fluxes. Therefore, the enzyme concentrations can be eliminated from the formulae. From the enzyme demand function Eq. (1), we obtain the function **z**(**x**), and inserting this into Eq. (5), we obtain the three score functions as functions of **x** alone:

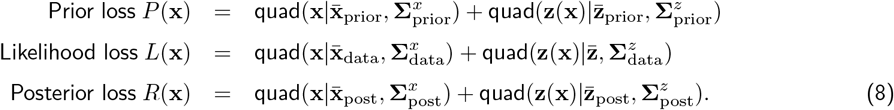

For distinction, in contrast to these “score” functions, the previous functions with ^*^ will also be called “prescores”. Our estimation method can be extended to problems with several metabolic states. Each condition *s* has its own flux distribution, metabolite data, and enzyme data, and we can run the estimation separately for each state (see also appendix A.1). In problems with several states, vanishing fluxes can be considered (see appendix B.3). In estimation problems with a single metabolic state, this is even simpler: reactions with vanishing flux can be omitted.

### 2.4 A convex version of the score functions

If an optimisation problems is convex, this has many advantages: it excludes separate local optima, decreases the numerical effort, and thus makes large problems tractable. Methods like parameter balancing and ECM profit from this. Are the prior, likelihood, and posterior score functions (8) convex in **x**? The first term in each score is convex because *F* is quadratic and therefore convex^11^. The second term is composed of a convex function quad(·) and a convex function **z**(**x**), but this alone does not make it convex. To be convex, it would have to be composed of an outer *non-decreasing* function and an inner convex function, but quad(*·*) is quadratic and obviously not non-decreasing.

However, here we argue that we can modify the second term to make it more similar to a convex function, in order to make the estimation problem better tractable. First, we can assume an uncorrelated prior for enzyme concentrations, i.e. a separate Gaussian prior for each reaction *l* and each metabolic state *s*. If we do this, the second term becomes a sum of the form

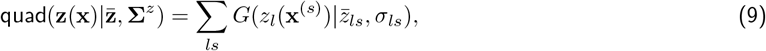

with a one-dimensional quadratic function 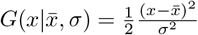. Next, we split this function into *G* = *G* _+_+ *G*_*−*_, where *G*_+_ = *G* only if *x* ≥ *a* (and 0 otherwise) and *G*_−_ = *G* only if *x* < *a* (and 0 otherwise). *G*_+_ is an increasing function, and the resulting truncated quadratic function

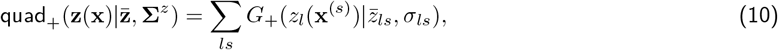

is a sum of non-decreasing functions *G*_+_(·) of a convex function **z**(**x**) and therefore convex (remember that the enzyme log demand ln **e**(**x**) is a convex function on the metabolite polytope [39] for a wide range of plausible rate laws.). If we now replace quad(·) by quad_+_(·) in all second terms of Eqs (8), we obtain the truncated score functions

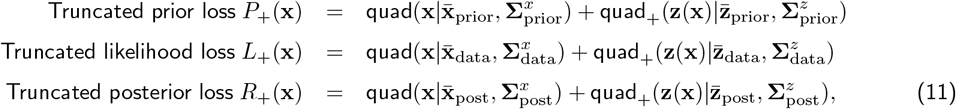

which are convex (or strictly convex if non-flat priors are used).

Now that the *G*_−_ term is gone, the posterior mode can be found by convex optimisation, while using the *G*_−_ term makes the problem non-convex^12^. Alternatively, we can replace *G*(·) by a relaxed version *G*_*α*_(·) = *G*_+_(·)+*α G*_−_(*·*), with a relaxation parameter *α ∈* [0, 1], which we can also see as an interpolation *G*_*α*_(·) = *α G*(*·*) + (1 − *α*) *G*_0_(·) between the original function *G* (if *α* = 1) and the truncated function *G*_0_ (if *α* = 0). Accordingly, we defined the relaxed score functions as interpolations

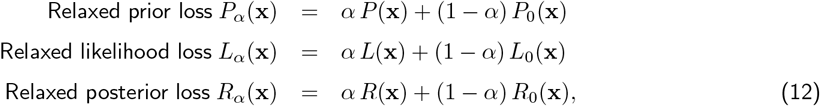

These functions are not convex (because they contain non-convex terms stemming from *G*_−_(·)), but by choosing small *α* values, they can be made arbitrarily similar to the convex truncated functions. By varying *α* between 0 and 1, we can interpolate between the convex truncated problem and the non-convex, original problem.

If we omit the *G*_−_ term, what will the effect on the estimation problem be? If we see *G* (in the terms related to enzymes) as a sum of *G*_+_ and *G*_−_, then *G*_+_ penalises enzyme concentrations that are higher than the posterior mean *z*_post_, while *G*_−_ penalises enzyme concentrations that are lower. If we omit the terms with *G*_−_, an estimated enzyme concentrations may be much smaller than the corresponding enzyme prior means or enzyme data, as if all this information did not exist. Thus, at this point, we have two possibilities: using the full posterior score function *R* (using the quadratic function *G*), which is non-convex, or using a posterior score function *R*_0_ without the *G*_−_ term (i.e., replacing *G* by *G*_+_) and accepting that enzyme concentrations may be underestimated. In fact, due to the relatively general form of our model, we expect that this trade-off (lower bounds or lower penalty terms for enzyme concentrations lead to non-convex estimation problems) holds for metabolic estimation problems more generally. However, we can still interpolate between the two possibilities. With *α >* 0, the relaxed posterior score function *R*_*α*_ will be non-convex, but by choosing small *α* values, we can make it arbitrarily similar to a convex one, while keeping at least a (weak) term for penalising small enzyme concentrations. Our tests (see below) show that this compromise, and the resulting estimation method is a promising tool.

### 2.5 Model balancing: estimation of kinetic constants and metabolic states

We now consider the full model balancing problem, that is, an estimation of kinetic constants, metabolite concentrations, and enzyme concentrations. Following [1], we parametrize the model by kinetic constants 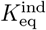, *K*_V_, *K*_M_, and possibly *K*_A_ and *K*_I_ (all in log-scale). For some of these constants, data may be available (for instance, equilibrium constants *K*_eq_ can be estimated from thermodynamic calculations) and consistent values need to be estimated. This problem resembles our simplified problem, where the enzyme concentrations were convex in **x**. Now the enzyme concentrations depend additionally on kinetic constants, and are convex in the joint space of log kinetic constants and metabolite log-concentrations (for any number of states)! This means that all the formulae and arguments for our previous simplified problem (including the usage of a “one-sided” enzyme posterior for obtaining a convex problem) can be reused. A description of the algorithm, including the convexity proof for the function ln *e*(ln **p**, ln **c**), is given in appendix A.2.

Since state variables and kinetic constants are estimated together, and since the kinetic constants are kept constant across metabolic states, the state variables become coupled across metabolic states and need to be estimated in one go. Instead of a metabolite vector **x**, we consider a larger vector **y**, containing the metabolite log-concentrations for all metabolic states and the vector of log kinetic constants. Allowed ranges and thermodynamic constraints define a solution polytope for the vector **y**. The prior, likelihood, and posterior loss scores contain terms that depend on enzyme concentrations **e**(**x**). If we insert Eq. (1) into these formulae, these terms are convex in the log kinetic constants, and independent of the metabolite concentrations^13^. Since *e*_*l*_(**q, x**) is a convex function of the vector 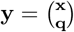, all terms of the likelihood loss score are convex in **y**. The prior loss score is strictly convex in **y** if a correlated prior for kinetic constants with pseudo values^14^ is considered [15]. Altogether, model balancing has the same favourable properties as the previous, simplified problem. In practice, the model balancing algorithm can be improved by a number of simplifications and tricks (appendix B). For example, enzyme concentrations (and therefore the likelihood function) increase very steeply close to thermodynamics-related polytope boundaries; to avoid numerical problems, a region close to the boundary may be excluded by extra constraints, or the log(log posterior) may be minimised instead of the log posterior.

## 3 Example application: Escherichia coli central metabolism

As a test case for model balancing, we consider a model of *E. coli* central metabolism taken from [39] (Figure 12 in appendix 12), and corresponding metabolite, enzyme, and kinetic data. We consider two different estimation scenarios, one with artificial data and one with experimental data from a single metabolic state (data from [39]). The algorithm was always run with the same settings (including priors and bounds). In all cases, we used the relaxed non-convex version of model balancing with a relaxation parameter *α* = 0.5. The calculation time (matlab function fmincon, with `interior-point` algorithm on a normal laptop) ranged between 2 and 30 minutes, with a single outlier of 150 minutes (notably, the numerical effort did not correlate with the size or expected difficulty of the problem, which suggests that the numerics could still be improved). So far, we did not test multiple realisations of the artificial data, and did not run the simulationswith multiple starting points. For details on model structure, kinetic and state data, and priors see appendix C.

**Figure 12.**
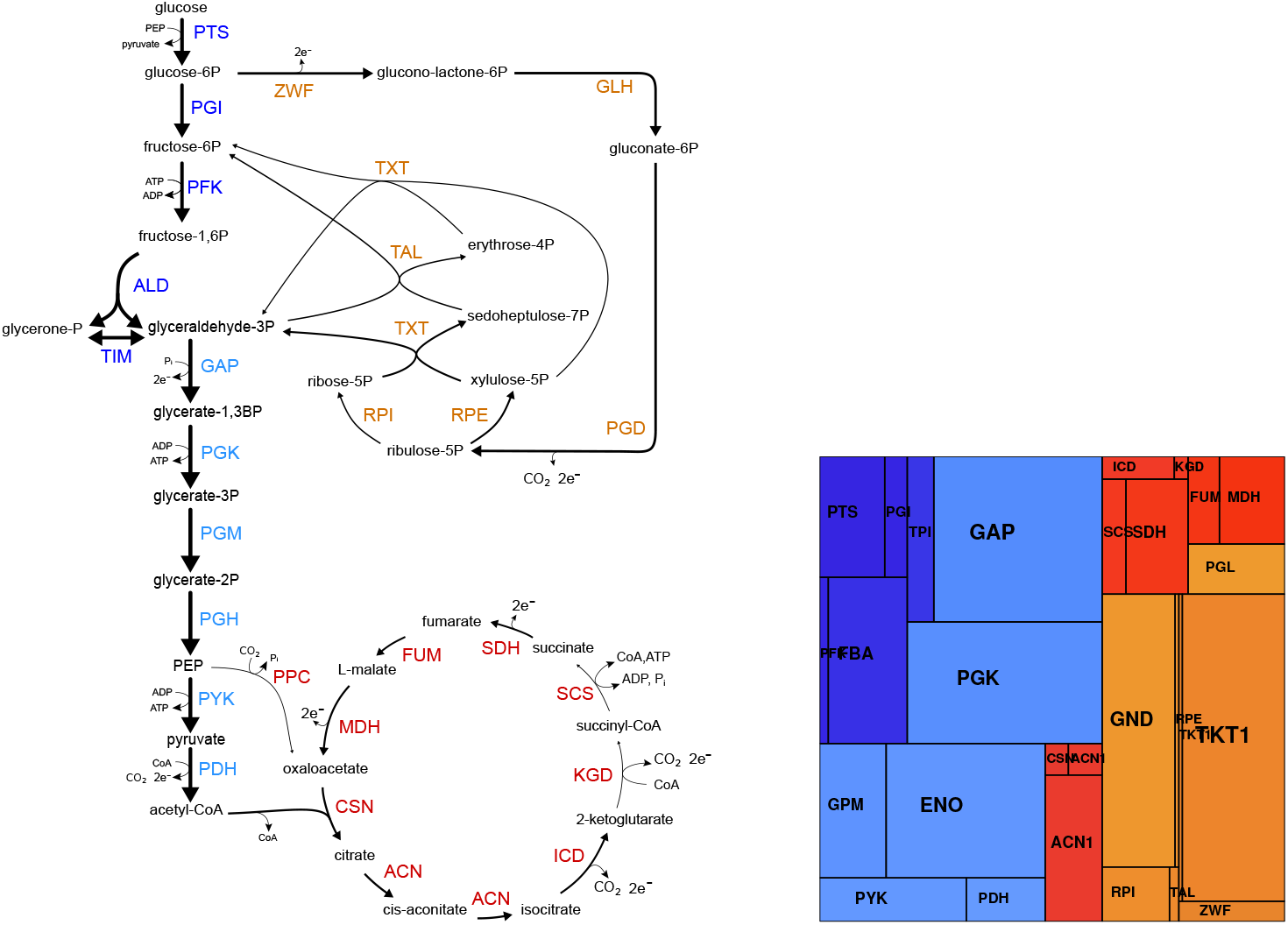
Model of *E. coli* central carbon metabolism and protein data, taken from [39].

### 3.1 Tests with artificial data

For the tests with artificial data, we first generated an artificial “true” set of kinetic constants and state data (metabolite concentrations, enzyme concentrations, and fluxes). Artificial measurement data were generated from the same random distributions that were also used as priors in model balancing. Metabolic state variables were generated from the kinetic model (with the “true” artificial kinetic constants) by randomly chosen enzyme concentrations and external metabolite concentrations and computing the steady states. For details see appendix C.2. Now the aim was to reconstruct the true (noisy-free) values for six simulated metabolic states from (noise-free or noisy) artificial data. We considered different scenarios (see Figure 13 in appendix) in which data were either fitted (metabolite and enzyme concentrations, and “known” kinetic constants) or predicted based on the other data (“unknown” kinetic constants). In the tests with artificial data, realistic statistical distributions (for kinetic constants, metabolic variables, and their measurement errors) were used to generate data, and the same distributions were used as priors when reconstructing the true values. To obtain realistic distributions of kinetic constants, we started from known (or suspected) distributions (from [39, 18], which relied on [12]), and adjusted them based on data. This is an ideal situation, making reconstruction particularly easy. In real-life applications, if our priors and assumed noise concentrations are not realistic, the reconstruction would be worse than suggested by our tests with artificial data.

**Figure 13.**
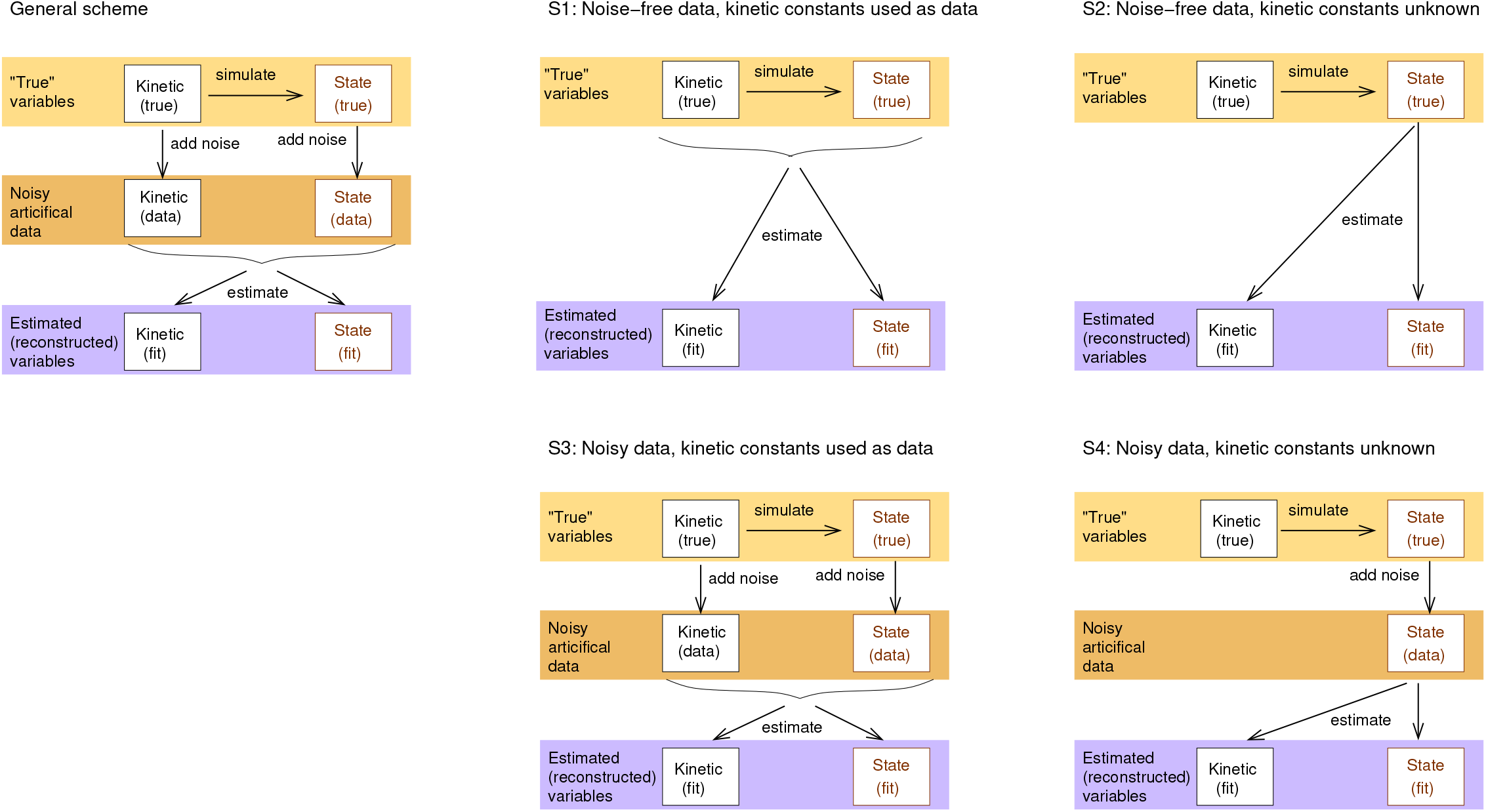
Estimation scenarios with artificial data. Left: general procedure. In a given model, kinetic constants are drawn from random distributions (respecting their interdependencies), and metabolic state data are generated by combining sampling and simulation runs (top row). From these “true” values, artificial (kinetic and state) data are generated by adding uncorrelated noise (centre row). Model balancing is used to estimate the kinetic parameters and metabolic state variables (bottom row), aimed to resemble the true values. Right: I employed four variants of this procedure (called S1-S4) in which noise is either considered or not (in the latter case, the noise level is set to zero), and kinetic data are used or not. In another variant, data for equilibrium constants are used as the only kinetic data.

The model balancing results with artificial data are shown in Figures 3, 4, 5, and 6, where noise-free or noisy kinetic and noisy state data were used in different combinations. In Figure 3 both kinetic and state data are free of noise. Subfigures show the different simulation and estimation scenarios (rows) and different types of variables (columns). Each subfigure shows a scatter plot between true and fitted variables (metabolite concentrations, enzyme concentrations, and different types of kinetic constants). Deviations from the diagonal indicate estimation errors. In the top row, data for all kinetic constants were used; in the centre row, only data for equilibrium constants were used, and in the bottom row, no kinetic data were used^15^ at all. Depending on the scenario, kinetic constants were either fitted (red dots) or predicted from data (magenta dots). The quality of fits or predictions is measured by geometric standard deviations^16^ and linear (Pearson) correlations (for logarithmic values). For comparison, we also estimated *k*_cat_ values from the same artificial data, using the “max-*k*_app_” method from [20], (Figure 7).

**Figure 3.**
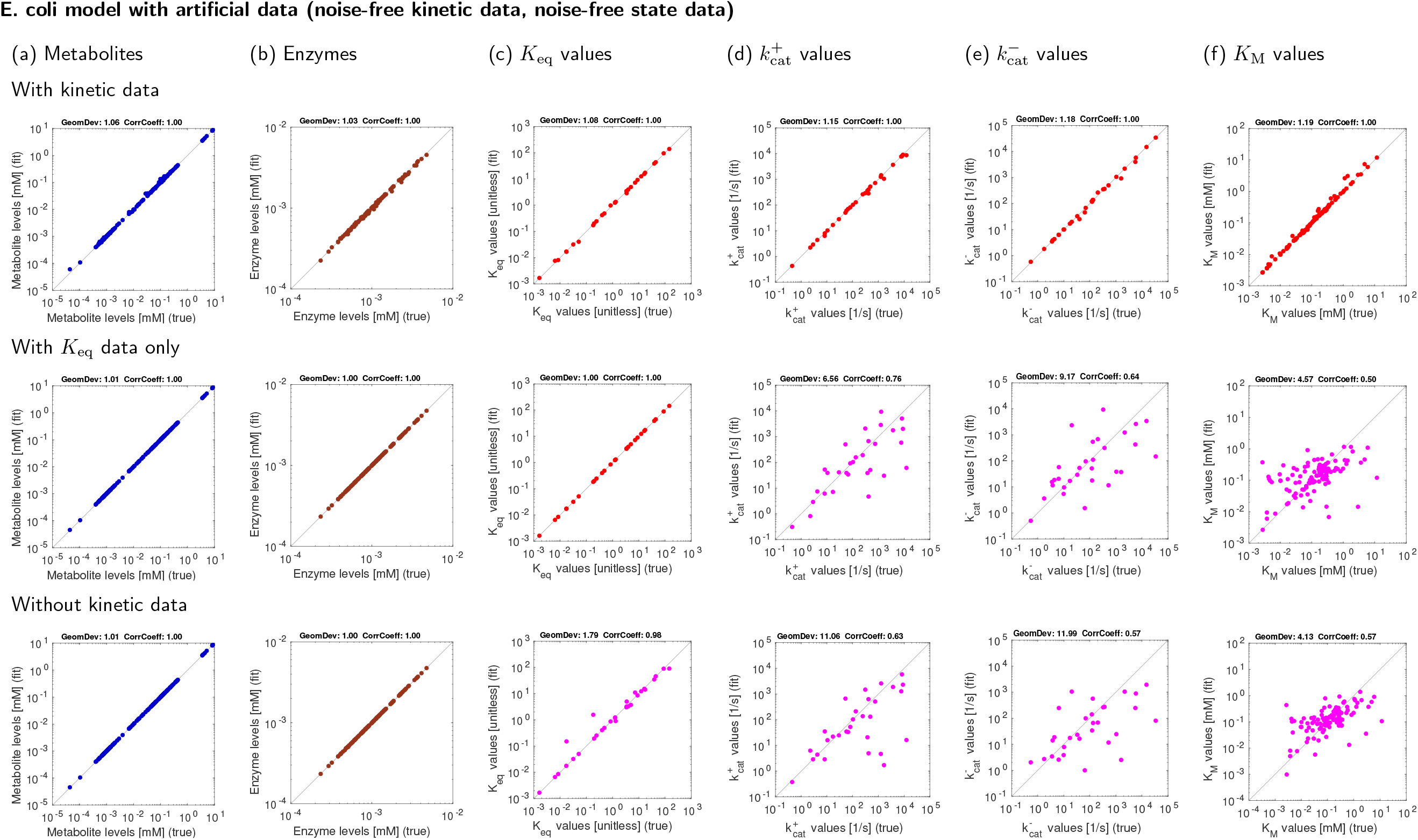
Model balancing results for *E. coli* central metabolism model with artificial data. The model structure is shown in Figure 12. Each subfigure shows “true” values (x-axis) versus reconstructed values (y-axis). Similarities are quntified by geometric standard deviations (“GeomDev”) and Pearson correlation coefficients (“CorrCoeff”). (a) Metabolite levels. (b) Enzyme levels. (c)-(f) Different types of kinetic constants. Rows show different estimation scenarios (see Figure) Upper row: simple scenario S1 (noise-free artificial data, data for kinetic constants). Centre row: scenario S1K (noise-free artificial data, kinetic data given only for equilibrium constants). Lower row: scenario S2 (noise-free artificial data, no data for kinetic constants). Depending on the scenario, kinetic constants are either fitted (red dots) or predicted (magenta dots).

**Figure 4.**
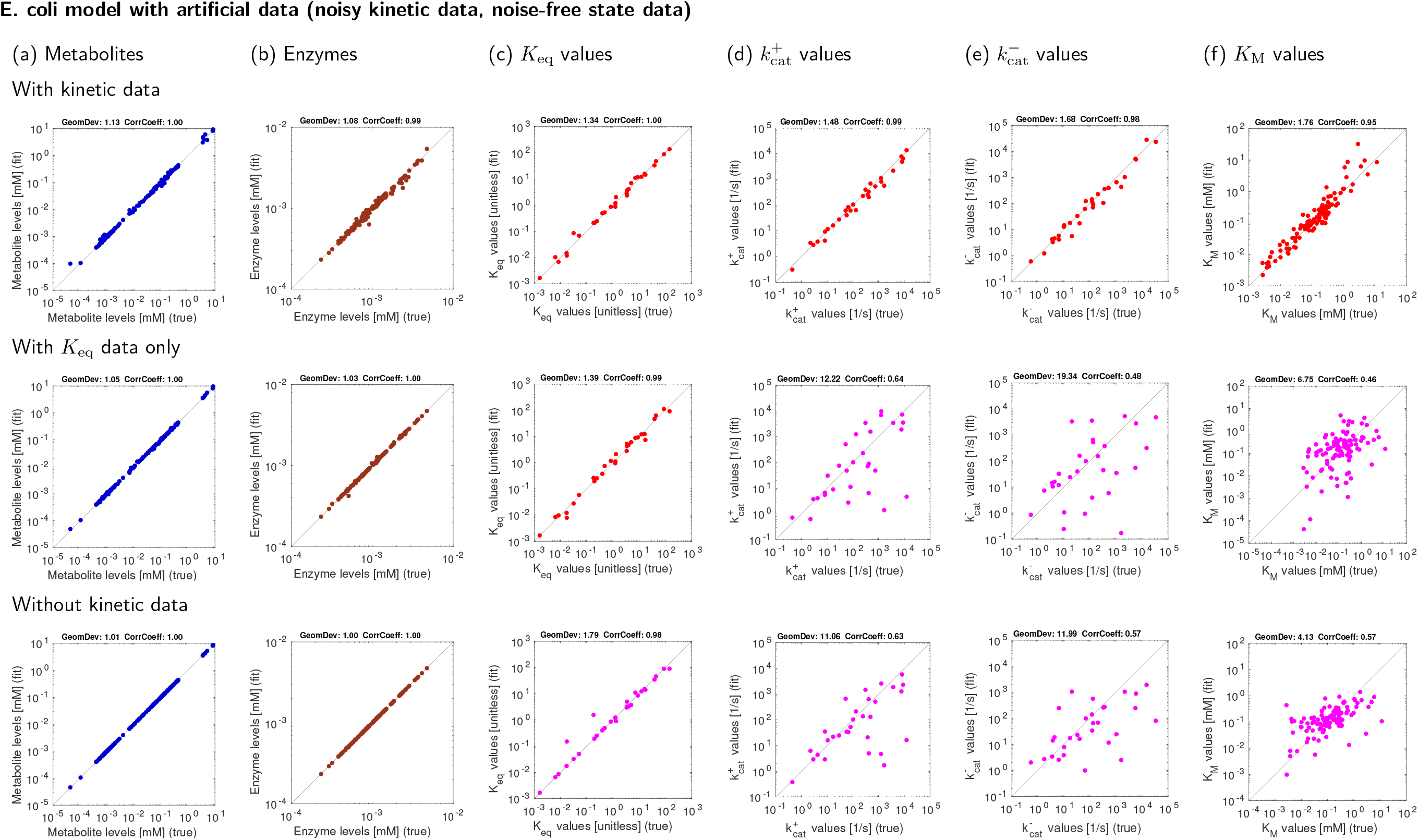
Same as Figure 3, with noisy kinetic data

**Figure 5.**
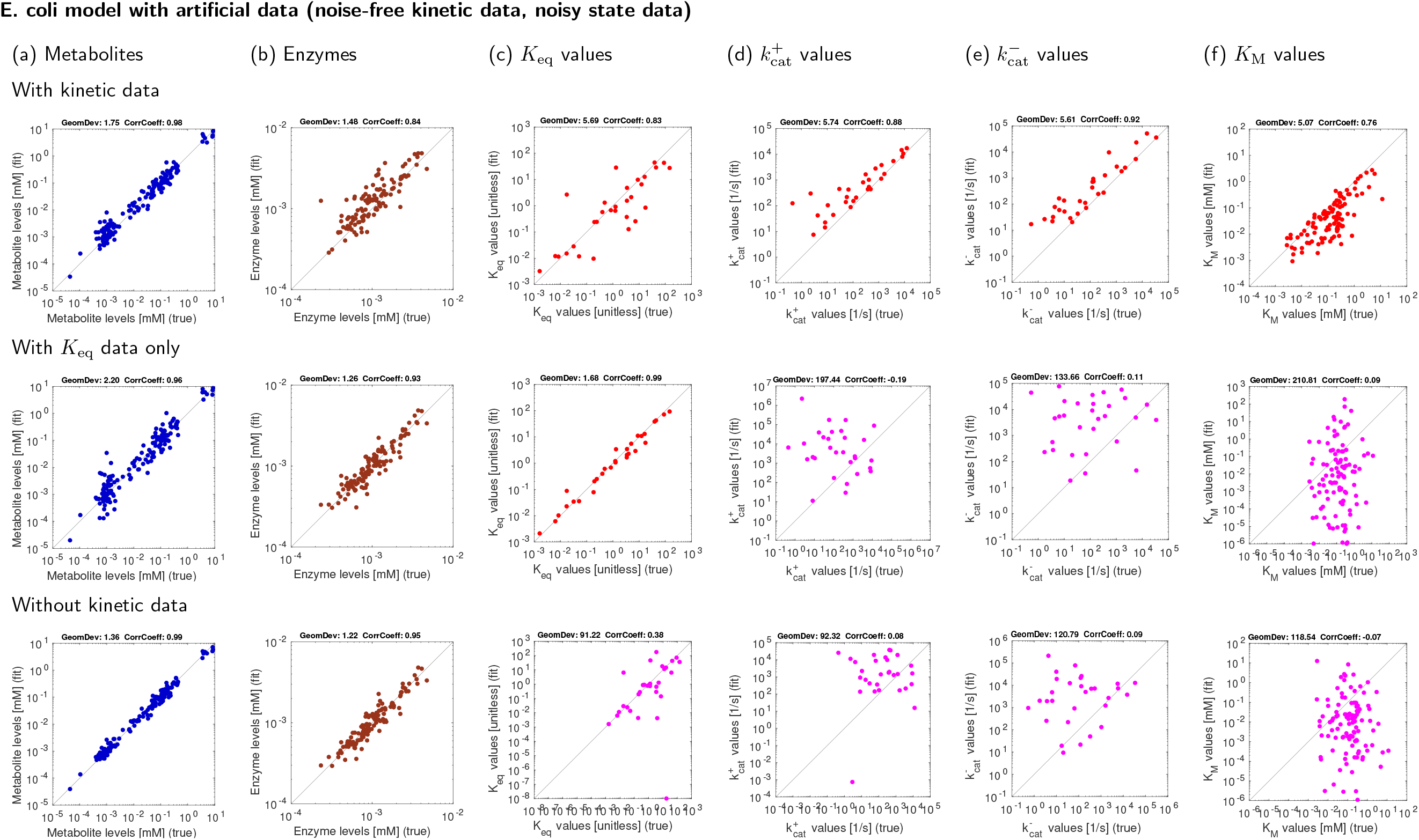
Results for *E. coli* central metabolism with noisy artificial data. Top row: estimation scenario S3 (noisy artificial data, data used for kinetic constants). Centre row: estimation scenario S3K (noisy artificial data, data for equilibrium constants only). Bottom row: estimation scenario S4 (noisy artificial data, no data for kinetic constants).

**Figure 6.**
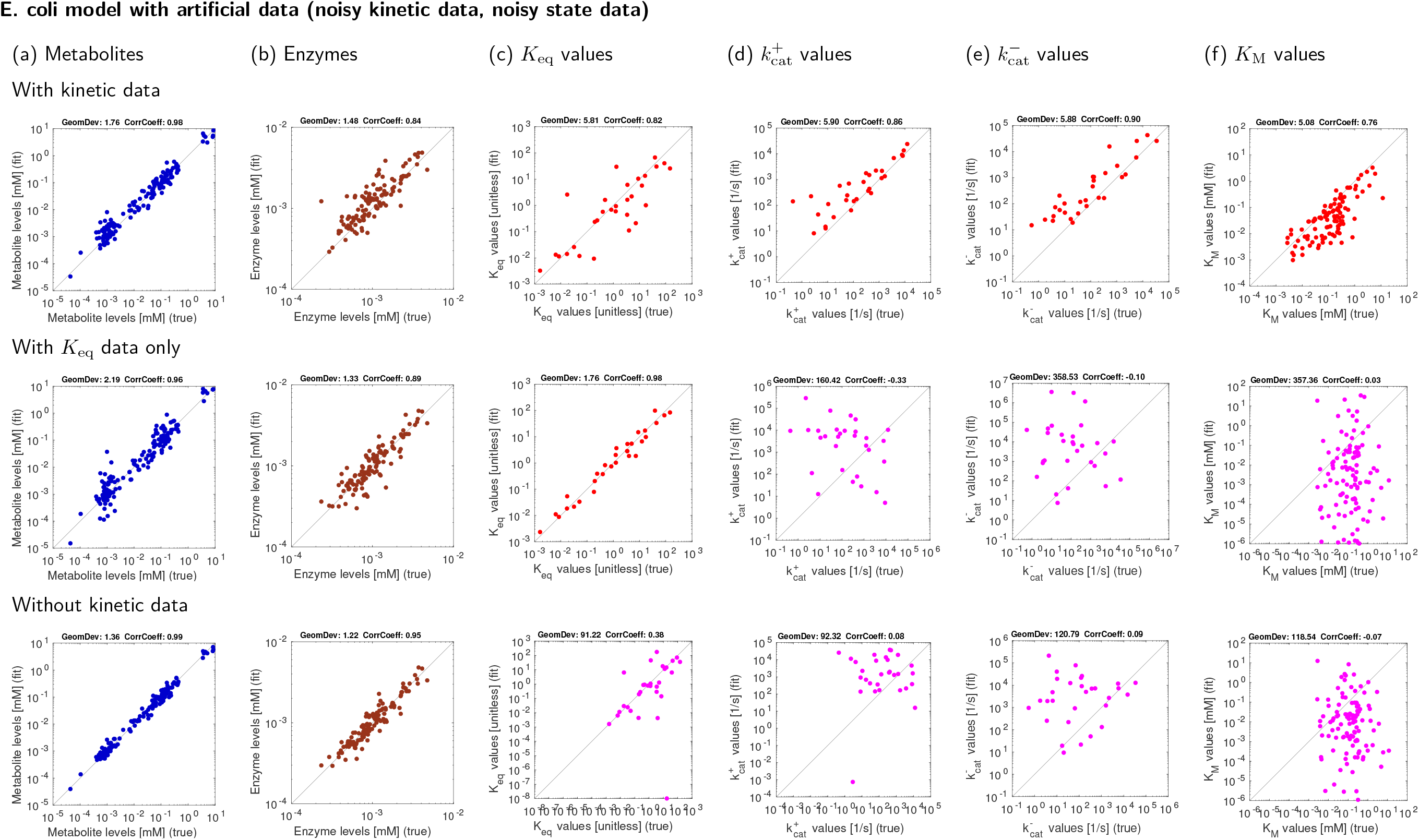
Same as Figure 5, with noisy kinetic data

**Figure 7.**
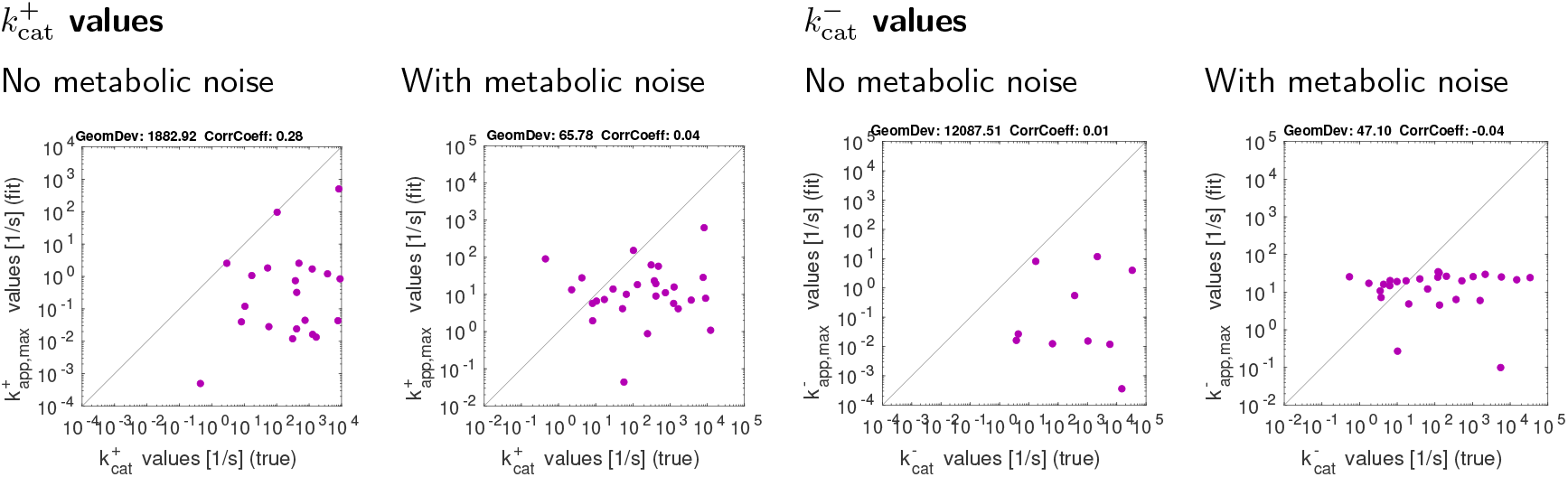
Catalytic constants in *E. coli* central metabolism (artificial data), estimated by kinetic profiling [20]. Note that *k*_cat_ values can only be estimated in the direction of fluxes (e.g. 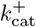 for reactions with forward fluxe).

The first scenario (top row) assumes ideal conditions: we assume noise-free, complete kinetic data and state data. Not surprisingly, the reconstruction errors are tiny, arising from small conflicts between data and priors. The other rows show estimation results using metabolic state data and equilibrium constants only (centre row) or using metabolic state data but no kinetic data at all (bottom row). The reconstructions in these two rows recover *k*_cat_ and *K*_M_ values partially, but they still depend on complete and precise state data. To assess the effect of noisy state data, we generated artificial state data (metabolite concentrations, enzyme concentrations, and fluxes) with a relative noise concentration of 20 percent. With noisy kinetic and/or state data, the estimation results become worse (Figures 4, 5, and 6), and especially the reconstruction of *K*_M_ values becomes very poor. Using data on equilibrium constants improves the results and *k*_cat_ values can still be partially reconstructed (Figure 6). The tests with artificial data show, first, that model balancing can adjust noisy data sets: relatively small changes in the data suffice to obtain a consistent set of kinetic and state data. Second, they show that information about *k*_cat_ values can be extracted from state data. Notably, even in the case without any kinetic data (nor equilibrium constants), model balancing yields better *k*_cat_ estimates than the kinetic profiling method. If we compare the results between Figures 3, 5 (last row, no use of any kinetic data) for model balancing and Figure 7 for kinetic profiling, the model balancing predictions are much more accurate in their indidivual values (Pearson correlation of 0.63 vs 0.28) if noise-free metabolic data are given. With noisy metabolic data, both methods show similar results. It turns out that *K*_M_ values are much harder to reconstruct: with noise-free metabolic data, a Pearson correlation of 0.5 (and a geometric standard deviation of 4.5) is achieved. With noisy metabolic data, the results are much worse: due to our priors, the estimates are in realistic ranges, but otherwise they appear to be randomly distributed. Therefore, with noisy metabolic data, adjusting in-vitro kinetic constants to in-vivo values (instead of estimating in-vivo values from omics data alone) remains the main application.

### 3.2 Tests with measurement data

After this test, we balanced the same *E. coli* model with real measurement data. As kinetic data, we used the *in-vitro* kinetic constants collected in [39] (“original kinetic data”) and, for comparison, a balanced version of the same data set (“balanced kinetic data”). For details on model and data, see appendix C.

Figures 8 and 9 show estimation results for a single metabolic state, aerobic growth on glucose (appendix C). Since the true model variables are not known, the available data are plotted against the reconstructed kinetic constants, metabolite concentrations and enzyme concentrations. In most cases, the scatter plots show the goodness of fit. In some cases, where some data were not used for fitting, the plots show the quality of actual predictions. In a first test, we used a consistent set of kinetic data obtained by parameter balancing as the kinetic data (Figure 8). If all kinetic data are used (top row), a good fit to these data is achieved, and only slight adjustments were necessary to obtain a consistent kinetic model agreeing with all the state data. However, we should not forget that the kinetic constants were fitted and not predicted (as indicated by red dots). In the centre row, equilibrium constants were used as the only kinetic data the other kinetic constants were actually predicted (as indicated by magenta dots). In the bottom row, all kinetic constants, including equilibrium constants, were predicted^17^. Now the correlations to *in-vitro* values are relatively small. This test may still be criticised because the kinetic data were taken from a previous parameter balancing procedure on the same network model and with the same priors, which might make it easier for model balancing to recover these values. To avoid any bias, we next ran model balancing with the original *in-vitro* kinetic data (which contain much fewer data points for comparison), with similar results.

**Figure 8.**
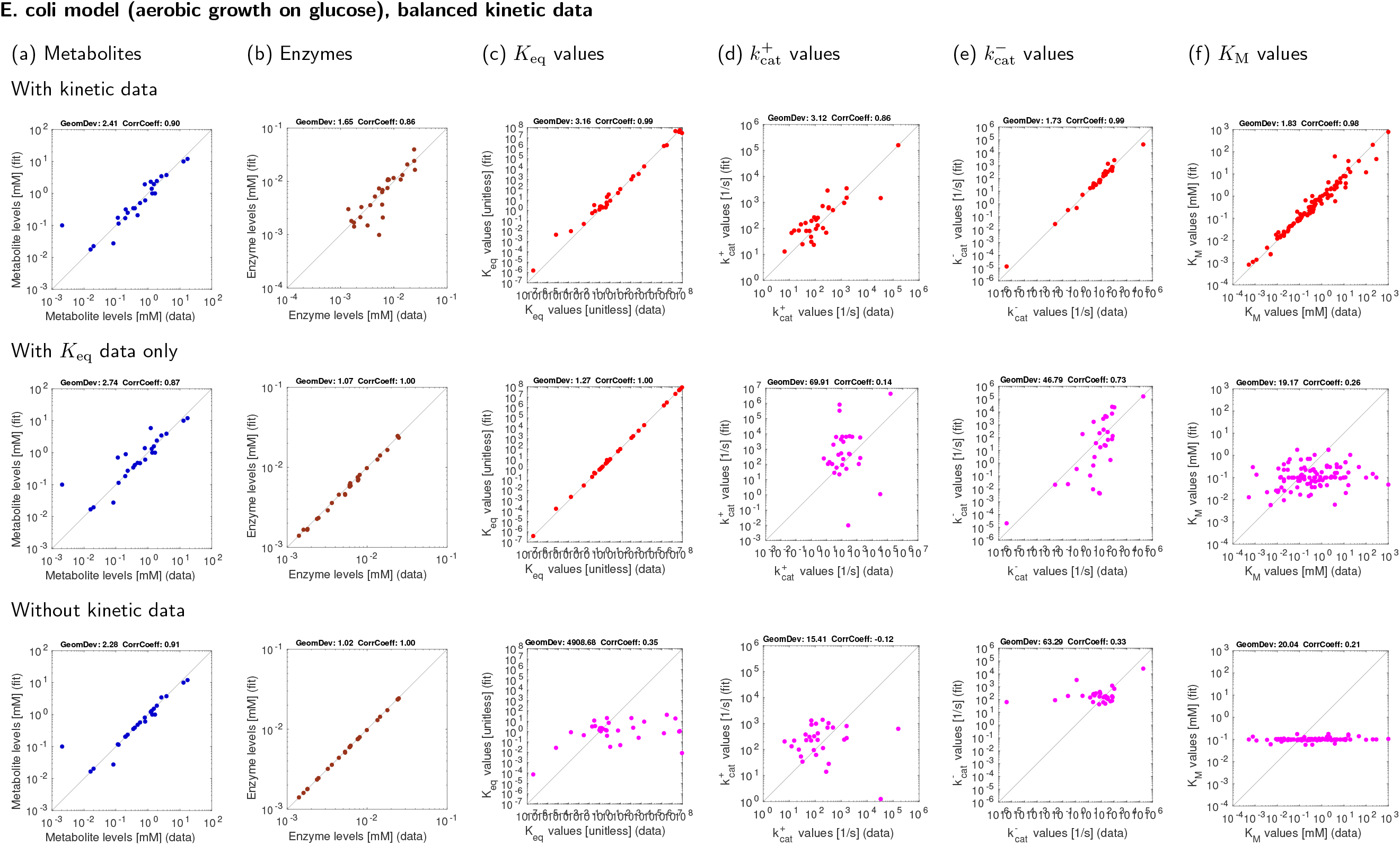
Results for *E. coli* central metabolism with experimental data (aerobic growth on glucose). The kinetic data stem from previous parameter balancing based on *in-vitro* data. Top: estimation using kinetic data. Centre: estimation using equilibrium constants as the only kinetic data. Centre: estimation using equilibrium constants as the only kinetic data. Bottom: estimation without usage of kinetic data. The same metabolite, enzyme, and kinetic data were used in [39].

**Figure 9.**
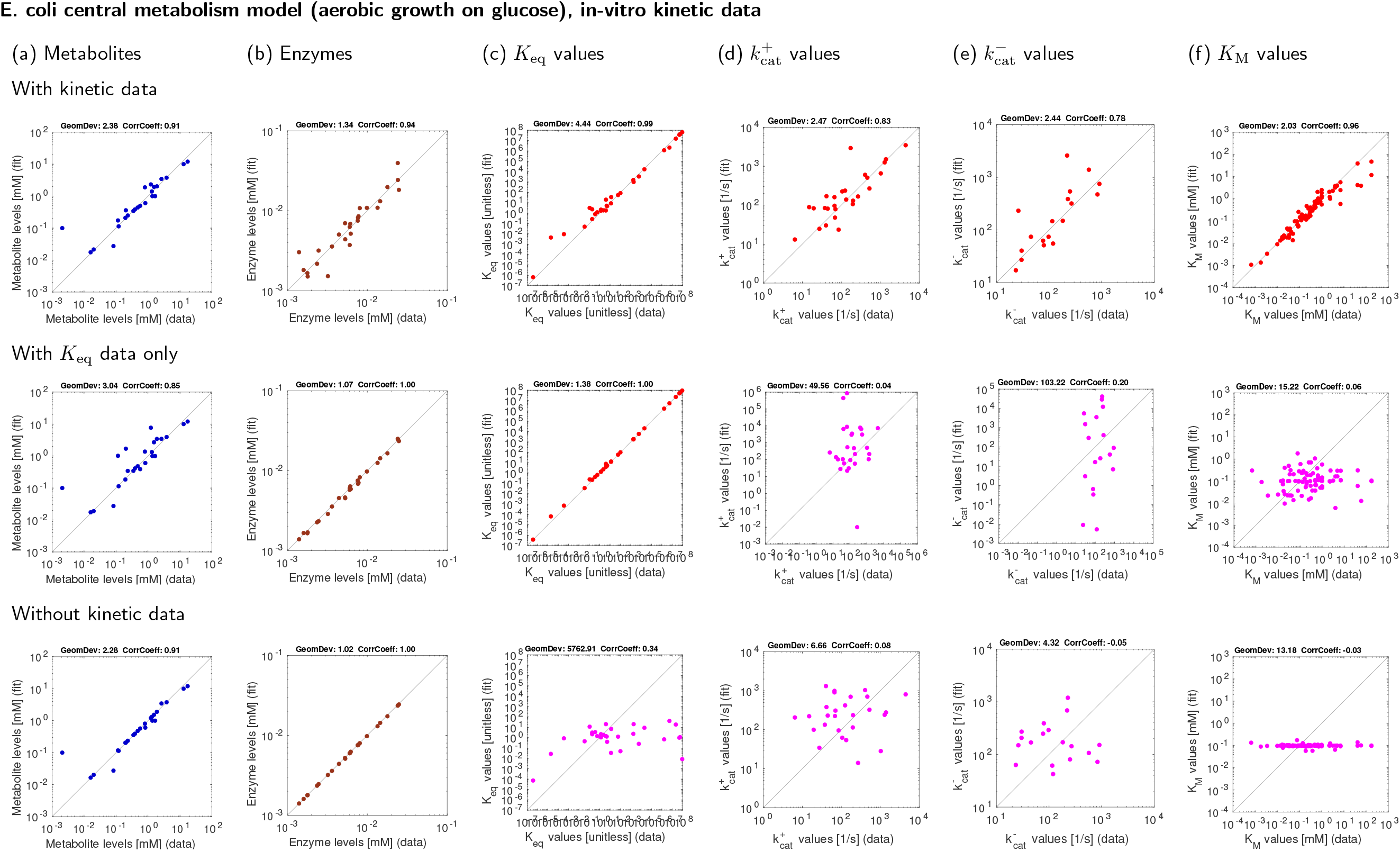
Results for *E. coli* central metabolism with experimental data (aerobic growth on glucose). Same as Figure 8, but based on original kinetic *in-vitro* data instead of balanced kinetic data.

**Figure 10.**
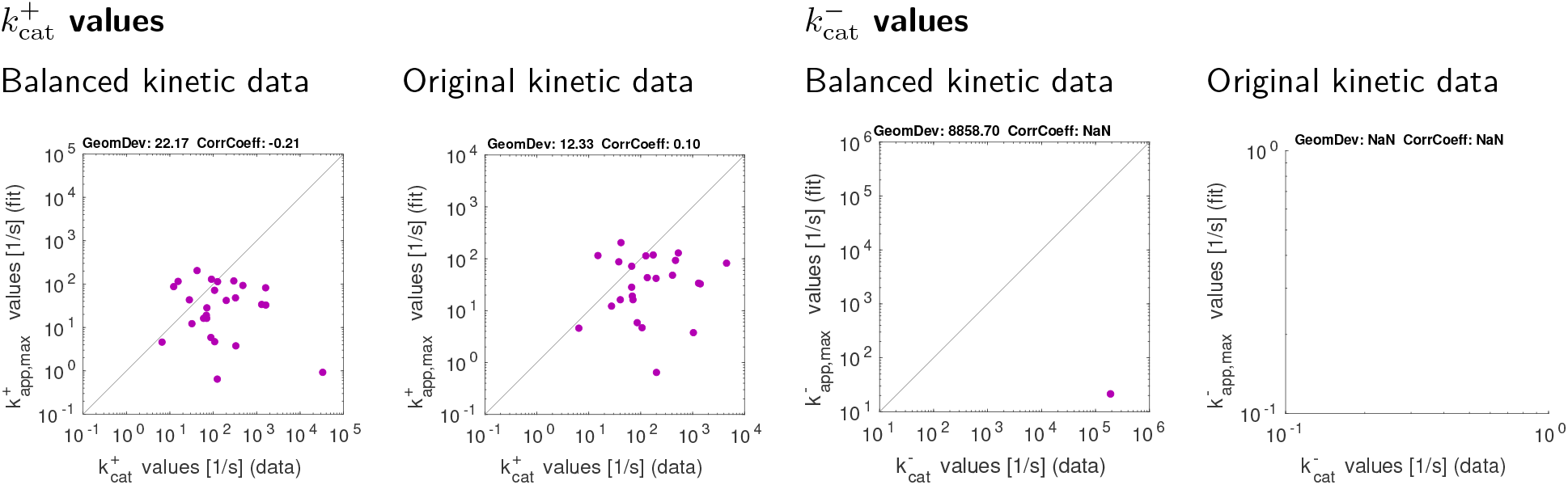
Catalytic constants in *E. coli* central metabolism model (aerobic growth on glucose), estimated by kinetic profiling [20].

### 3.3 Parameter identifiability

To understand how much useful information can be extracted from (noise-free or noisy) data, we need to think about uncertainties, parameter identifiability, and the usage of priors. In model fitting, generally, certain parameters or parameter combinations may be non-identifiable, that is, they have no effects on measurable outcomes, and therefore we cannot determine them from available data [24]. Likewise, if parameters have very small effects on measurable outcomes, and if data are sparse or noisy, our estimates of these parameters will be very uncertain. For example, if kinetic constants are estimated from metabolic state data, and if a reaction is strongly forward-driven, its backward *k*_cat_ and *K*_M_ values may be hard to determine (compared to forward *k*_cat_ and *K*_M_ values value); if an enzyme is always close to saturation, its *K*_M_ value will be poorly determined, and if an enzyme remains always in its linear range (due to small substrate levels), only the *k*_cat_*/K*_M_ ratio will be determined, but not *k*_cat_ and *K*_M_ individually; finally, enzyme concentrations and *k*_cat_ values appear in rate laws in the form of a product (which is therefore well-determined by the fluxes), but their ratios may be ill-determined. In particular, an error in the scaling of flux or enzyme data will have an immediate effect on *k*_cat_ estimates.

For model balancing, this means: variables or variable combinations (e.g. *k*_cat_*/K*_M_ ratios) may be structurally non-identifiable (if they cannot have any effect on the metabolic state), practically non-identifiable (for example, if data from *n* states would be needed to infer them, but less data are given), or poorly identifiable (if they have a very small effect, e.g. the *K*_M_ value in an enzyme close to full saturation). Since model balancing also uses priors and kinetic data, non-identifiability does not mean that parameters are ill-determined: the method will always result in a posterior with clearly defined modes – however, in these solutions the values of “non-identifiable” kinetic constants or state variables will completely depend on the priors, and will therefore be sensitive to the (somehow arbitrary) choice of the prior mean. Non-identifiability will always arise if there are fewer data values than variables to be estimated. For example, state data from a single metabolic state may not suffice to reconstruct the kinetic constants; if more metabolic states are used, the kinetic constant may become well-defined.

If model variables are non-identifiable (or poorly identifiable), this will lead to considerable uncertainties, visible in the posterior distribution. Assuming that the problem is convex, and that non-flat priors are used, we know that the posterior mode is single point in solution space. However, if the priors are broad, this point is somewhat arbitrary, and posterior sampling would reveal that many other solutions (along the “non-identifiable subspace”) are almost as good, revealing a large uncertainty. This raises some practical questions. Can we improve the result by using more data (e.g., metabolite concentrations from more metabolic states)? If no kinetic data are given, how many metabolic states are needed to identify – at least potentially – all kinetic parameters? Which variables are hard to reconstruct? And are there variables that remain non-identifiable, no matter how much state data we use? Paradoxically, variables that are *always* non-identifiable (or poorly identifiable) are less of a problem, because these variables, in turn, are also irrelevant for model predictions. An example would be an enzyme that is *always* close to saturation – we cannot determine its *K*_M_ value, but we also do not need this *K*_M_ value for forward simulation. However, if we try to simulate other conditions, in which the enzyme *is not* close to saturation, the results may be poor. Of course, the identifiability problem is not specific to model balancing; other estimation methods would face the same problem.

## 4 Discussion

Model balancing completes the earlier attempt from [14] to integrate kinetic and metabolic data to obtain a consistent metabolic model with several states within a Bayesian framework, accounting for uncertainties and missing data. Depending on data availability and quality, it can be used for (i) inferring kinetic constants from state data and (ii) making almost complete, uncertain data consistent, certain, and precise. Various methods and tools have previously been developed to parameterise metabolic models, to integrate omics data, and to build model ensembles. These methods use different types of knowledge (*in-vitro* kinetic constants, omics data, and physical parameter constraints) and different estimation approaches (including machine learning, regression models, calculations based on rate laws, and model fitting).

1. **Ensemble modelling** If the parameters or states of a model are not described by fixed numbers, but by probability distributions, we obtain a model ensemble. Ensemble modelling can be used to express the actual possibilities of a system or the possibilities we attribute to it given our limited data and knowledge, and may concern not only possible steady states, but also dynamic behaviour. Structural Kinetic Modelling (SKM) [34], for example, is a method for model parameterisation in which parameters are not fitted but chosen randomly to create a model ensemble. A consistent model state is constructed in two steps: first, a metabolic state is defined by choosing fluxes and metabolite concentrations. Then, kinetic constants are chosen at random in agreement with the predefined metabolic state. In practice, this is achieved by randomly sampling the saturation values of enzymes and then reconstructing the corresponding kinetic constants. Structural thermokinetic modelling (STM) [40], a variant of SKM, considers reversible rate laws and guarantees thermodynamically consistent results. The procedure follows a dependency schema just like in model balancing. In the first step, it requires thermodynamically consistent fluxes, metabolite concentrations, and thermodynamic forces. In the second step, thermodynamic forces are used to convert saturation values into correct reaction elasticities. SKM and STM can be adapted to account for priors or data for the *K*_M_ values. However, including data or priors for *k*_cat_ values and enzyme concentrations remains difficult, and the method cannot be used to used to fit kinetic constants simultaneously to several metabolic states.
2. **Parameter estimation** In parameter estimation, the idea of ensemble modelling is turned upside down. Instead of assuming a distribution of model parameters and doing forward simulations, we infer such a distribution from available data, reflecting our knowledge about the model and its parameters. In practice, parameter fitting in kinetic models can be performed by Monte-Carlo methods such as random screening, genetic algorithms, or simulated annealing. For example, one may generate a large ensemble of possible parameter sets, compute the likelihood or posterior density values for each of them, and choose the best parameter set among them (see [33] for an example). Sampling-based optimisation methods are generic and easy to implement, but with large parameter spaces and complicated objective functions, finding the optimum becomes very unlikely. Moreover, without understanding the optimality problem, it is hard to assess whether there are many local optima, and how good the computed solutions actually are.
3. **Fitting kinetic constants in single reactions: SIMMER and kinetic profiling** Estimating kinetic constants becomes much simpler if it can be done separately for different reactions. For example, if the flux, metabolite concentrations, and enzyme concentration in a reaction are known (completely and precisely) in several metabolic states, the kinetic constants can be fitted without knowing the rest of the system^18^ [44, 21]. However, this approach has a number of limitations: to estimate the kinetic constants of a reaction, all pieces of information, including the concentration profiles of all reactants, must be given. Model balancing, in contrast, attempts to reconstruct missing information from other parts of the network. Kinetic profiling, an approach to estimate *k*_cat_ values, has been described in the introduction. It is easy to implement, and – given protein concentrations and fluxes – the computational effort is low. However, it provides only a lower bound, which may be far from the true value if only few metabolic states are considered. Model balancing, in contrast, has the potential to integrate different types of data (including *K*_eq_ values and *in-vitro k*_cat_ values) to obtain a best estimate, and is in principle able to estimate a *k*_cat_ value even if this value is never reached in a living cell.

Parameter estimation in kinetic models can easily lead to non-convex optimality problems, especially if it involves a screening of thermodynamically feasible flux distributions [45]. To obtain problems that are nearly convex, model balancing relies on two insights: all fluxes must be predefined^19^, and log kinetic constants and metabolite concentration must be used as free variables^20^. Model balancing combines ideas from two previous, convex methods: Parameter Balancing (PB) for the estimation of kinetic constants (and possible metabolic states) from data, and Enzyme Cost Minimisation (ECM) for predicting biologically favourable states from a principle of minimal effort (see Figure 2).

**Figure 2.**
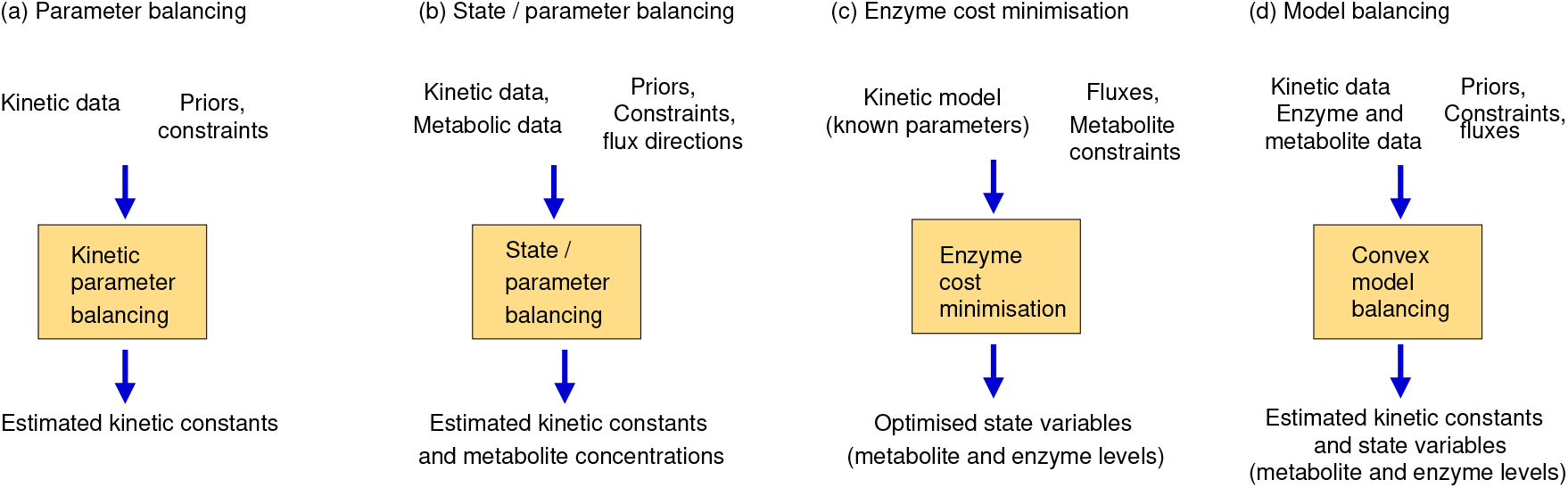
Model balancing and similar methods for parameter estimation and optimal metabolic states. The methods differ in their purpose (parameter estimation versus prediction of biologically optimal states), the choice of free variables (kinetic constants and/or metabolite and enzyme concentrations), and data used, but they all share some mathematical features: kinetic constants and metabolite concentrations are described in log-scale (such that all dependencies become linear); thermodynamic and physiological constraints are imposed; and fluxes are predefined. In each of these methods, the search space is a convex polytope. In the previous methods, the objective function was convex (either quadratic or derived from kinetics), leading to convex optimality problem. In model balancing, the non-convex terms (which penalise low enzyme concentrations) can be weakened or omitted to make the problem easily solvable.

1. **Parameter balancing**. Parameter balancing is a method for estimating consistent kinetic and thermodynamic constants from kinetic and thermodynamic data. It resembles model balancing, but (in its basic form) does not use detailed information on rate laws and fluxes. All multiplicative constants (such as Michaelis-Menten constants or catalytic constants) are described by log-values. To account for parameter dependencies^21^, we choose a subset of kinetic constants^22^, the free parameters in our linear regression model, and express all other kinetic constants as functions of them. With Gaussian priors and measurement errors (in log-scale), likelihood and posterior terms are quadratic and convex. Parameter balancing can also be applied to kinetic and thermodynamic constants (“kinetic parameter balancing”), to metabolite concentrations and thermodynamic forces in one or more metabolic states (“state balancing”), or to kinetic constants and metabolic states together (“state/parameter balancing”). Known flux directions can be used to define the signs of thermodynamic forces. Thus, parameter balancing can predict thermodynamically feasible kinetic constants and metabolite concentrations and its optimisation takes place on the same set as in model balancing. It provides plausible ranges for kinetic constants, but unlike model balancing it does not consider rate laws and cannot be used to fit kinetic constants to data^23^.
2. **Enzyme cost minimisation**. Enzyme cost minimisation (ECM) [39] predicts optimal enzyme and metabolite concentrations in kinetic models with given parameter values. ECM assumes predefined metabolic fluxes and determines metabolite and enzyme concentrations that realise these desired fluxes at a minimal cost, where cost functions can be a linear or convex function of the enzyme concentrations, plus a convex function of the metabolite concentrations. Thus, in contrast to parameter balancing, this method uses kinetic rate laws with given kinetic constants, and it is a biological cost (typically, a cost depending on absolute metabolite and enzyme concentrations), not a fit to data, that is optimised. The optimisation is carried out in (log-)metabolite space. With given rate laws, the enzyme concentrations can be written as functions of metabolite concentrations and fluxes and the cost function (a weighted sum of enzyme and metabolite concentrations) is convex on the feasible metabolite polytope.

Model balancing combines elements from these two methods. Like in parameter balancing, kinetic constants and metabolite concentrations are used as free variables (forming a parameter/concentration polytope as the solution space). And, like in ECM, we assume that the fluxes are given and use the fact that the (now, logarithmic) enzyme concentrations are convex functions of the metabolite log-concentrations. We combined this with two additional insights: the fact that (both absolute and logarithmic) enzyme concentrations are convex functions on the *combined solution space* of kinetic and metabolic variables, and the fact that most of the terms in the posterior score are convex, while the one sort of terms (limiting enzyme concentrations from below) that is non-convex can be omitted or be “weakened” by a tunable prefactor.

In all three methods – Parameter Balancing, Enzyme Cost Minimisation, and Model Balancing – the solution space is a high-dimensional polytope (in the space of log kinetic constants, metabolite log-concentrations, or both). To construct this polytope, we define a box by upper and lower bounds and add linear constraints describing dependencies. In fact, the solution polytope for Model Balancing is obtained from the polytopes of the other methods by taking their Cartesian product and removing infeasible regions, in which constraints between kinetic constants and metabolite concentrations would be violated (shown in Figure 11). Since all variables are estimated together, any information about one variable (bounds, priors, or data) can improve the estimates of other variables, too. In parameter balancing, a data value for a kinetic constant may improve the estimates of all other variables. Similarly, in model balancing additional metabolite and enzyme data may also improve the estimation of all kinetic constants.

**Figure 11.**
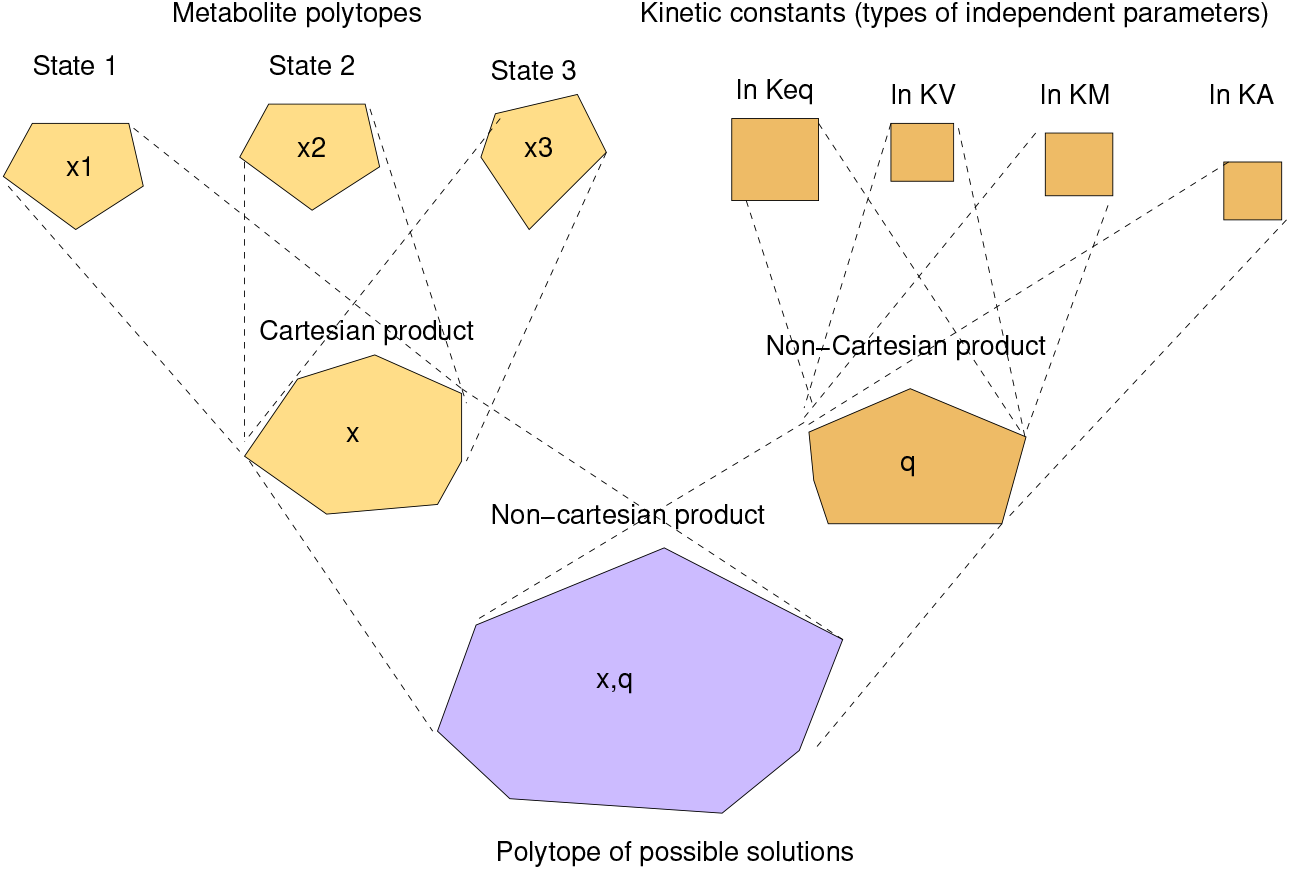
Solution space in model balancing. The free model variables (metabolite concentrations and kinetic constants, all on log-scale) are constrained by physiological bounds and thermodynamic constraints dependent on flux directions. Together, all these inequality constraints define a feasible region in the space of log-variables (bottom), called the solution space. This high-dimensional polytope arises from a “non-Cartesian” product between a metabolite polytope and a kinetic constant polytope (centre), a Cartesian product from which some parts are removed due to constraints. The metabolite polytope itself is a Cartesian product of the metabolite polytopes for single metabolic states (which may differ between each other if flux directions are different); the polytope of kinetic constant is a (non-Cartesian) product of polytopes (boxes) for the different types of kinetic constants (top).

In summary, model balancing can use various types of knowledge (network structure, data, priors, and constraints), handles different types of variables (as defined by the dependency schema used), and applies to steady and non-steady states^24^ as long as the fluxes are thermodynamically correct^25^. The tests with articifial and measurement data show that, in realistic scenarios, a consistent model with consistent metabolic states can be obtained by relatively small adjustements of the data, that precise metabolic state data and equilibrium constants contain information about *k*_cat_ and *K*_M_ values, but that the estimation of *k*_cat_ and *K*_M_ values from metabolic data with realistic noise levels is relatively poor (although it tends to be better than the prediction by the existing kinetic profiling method). We did not perform a systematic leave-one-out crossvalidation for individual kinetic constants, but the outcome of such a crossvalidation can be expected to lie in between our two scenarios in which either all kinetic constants were used as data, or only equilibrium constants were used as data. We can conclude for sure that the usage of equilibrium constants improves the results, which confirms the importance of known equilibrium constants for constructing reliable kinetic models.

Depending on the data available, model balancing can be applied in different ways.

1. **Infer missing data types** Sometimes, data for two of our data types (kinetic constants, metabolite concentrations, and enzyme concentrations) are available and the aim to estimate the third, missing type of data. There are three cases: we may estimate *in-vivo* kinetic constants from fluxes, metabolite concentrations, and enzyme concentrations; we may estimate metabolite concentrations from fluxes, enzyme concentrations, and a kinetic model; or we may estimate enzyme concentrations from fluxes, metabolite concentrations, and a kinetic model. If the data were complete and precise, the third type of variables could be directly computed without model balancing. But if data are uncertain and incomplete, model balancing allows us to infer the missing data while completing and adjusting the others.
2. **Adjusting omics data to obtain complete, consistent metabolic states** If the kinetic constants are known and metabolite and enzyme have been measured, we can turn these incomplete and uncertain data into consistent and plausible metabolic states, in agreement with our model. Again, fluxes must be given and their directions must agree with the assumed equilibrium constants and metabolite bounds. Even without any enzyme or metabolite data, we can still apply model balancing: in this case, the method would predict possible metabolic states with the given fluxes, relying on priors for enzyme or metabolite concentrations.
3. **Imposing thermodynamic constraints and bounds and data** To obtain a consistent model, we may collect data for kinetic and state variables and use model balancing to translate them into parameters and state variables. The resulting values will satisfy the rate laws, agree with physical and physiological constraints, and resemble the data and prior values. As in all other cases, posterior sampling could be used to decrease and assess uncertainties about the model parameters.

Model balancing extracts information from heterogeneous data. Even if almost no data are available, it can be used to obtain plausible models or model ensembles. In the tests with articificial data, model balancing performed well when precise data were given, and even with imprecise data it performed better than estimation by kinetic profiling. A main limitations of model balancing is that, unlike its predecessor methods, it is not a convex problem unless the *G*_*−*_ terms (which penalise low enzyme concentrations) are omitted. By downscaling these terms with a constant factor, the the posterior score can be made more similar to a convex function (which may make it easier to treat numerically), but it remains non-convex unless these terms are omitted completely. Another limitation of model balancing is its calculation speed. The number of variables increases with model size and the number of metabolic states, which impacts memory requirements and calculation time (results not shown). Due to its simplicity, kinetic profiling is much faster.

Here we considered a Bayesian estimation problem, constructed the posterior, but otherwise did not follow a Bayesian methodology. Instead of sampling from the posterior, we only considered the problem of finding the posterior mode. Posterior sampling would be big step forward, allowing us to determine the posterior mean, variance, covariances and marginal distributions of kinetic constants and state variables. Moreover, entire parameter sets could be sampled to obtain a model ensemble. To simplify sampling, the posterior may be approximated by a multivariate Gaussian distribution, obtained from the posterior mode and the Hessian matrix of the posterior score in this point. Such a sampling procedure has been implemented for parameter balancing, but not yet for model balancing. We hope that our formulation of the problem will facilitate posterior sampling, due to the fact that the posterior score is close to convex, and that it will pave the way to efficient sampling algorithms that might also include a sampling of metabolic fluxes [45].

## Acknowledgements

Many ideas for this work were developed in the European Commission 7th Framework project BaSysBio (LSHG-CT-2006-037469). We thank Marc Dinh and Vincent Fromion for insightful discussions about parameter identifiability.

## A The model balancing problem

### A.1 Model variables and constraints

To define a model balancing problem, we consider all kinetic constants and state variables (as “model variables”) and describe their dependencies. To do this, we split the model variables into “independent” (or “free”) variables and “dependent” variables based on the following thoughts. (i) To account for dependencies between kinetic constants, we treat some of them as free variables (independent equilibrium constants^26^, Michaelis-Menten constants, and velocity constants), while all others are dependent on them (dependent equilibrium constants, and catalytic constants). With kinetic constants described in log-scale, these are linear dependencies. (ii) For each metabolic state, we consider a metabolite log-concentration vector, an enzyme concentration vector, and a flux vector. Vectors from different metabolic states (usually given as columns of a data matrix) are merged into a large vector. (iii) Since enzyme concentrations depend on kinetic constants, metabolite concentrations, and fluxes, they are treated as dependent variables. (iv) Thermodynamic driving forces depend on equilibrium constants and metabolite concentrations, and are therefore dependent variables. Thus, the kinetic constants and metabolite concentrations remain the only free variables. (v) The predefined flux directions determine the signs of thermodynamic driving forces, which in turn imply linear constraints between equilibrium constants and metabolite concentrations on log-scale. Altogether, we obtain the following variables and dependencies (see Figure 1 (b)).

1. **Independent variables** Our free variables comprise (i) the independent kinetic constants in log-scale (in-dependent equilibrium constants 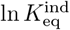, Michaelis-Menten constants ln *K*_M_, allosteric activation constants ln *K*_A_, allosteric inhibition constants ln *K*_I_, and velocity constants ln *K*_V_), collected in a vector

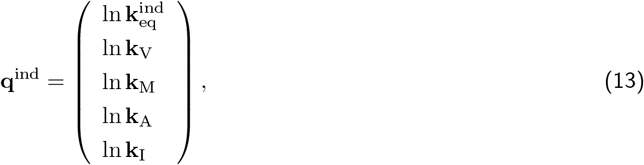

and (ii) the metabolite log-concentrations from one or several metabolic states *s*, contained in metabolite vectors **x**^(*s*)^ = ln **c**^(*s*)^. We obtain a vector of free variables

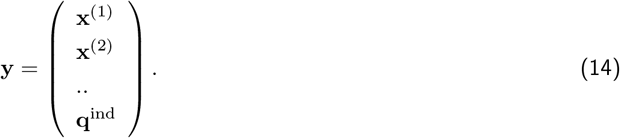

With *n*_*p*_ independent kinetic constants, *n*_*m*_ metabolites, and *n*_*s*_ metabolic states, the vector contains *n*_*p*_ + *n*_*m*_ *n*_*s*_ free variables.
2. **Dependent variables** We consider three types of dependent variables: dependent kinetic constants, enzyme concentrations, and thermodynamic forces. (i) The dependent kinetic constants on log-scale (dependent equilibrium constants 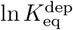, forward catalytic constants 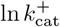), and backward catalytic constants 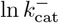) form a vector

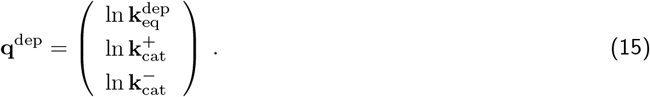

This vector can be computed from **q**^ind^ with the help of a linear function **q**^dep^ = **M**^dep^ **q**^ind^. The dependency matrix **M** follows from the stroichiometric matrix as described in [15]. Similarly, the vector **q** of all kinetic constants is given by a linear formula

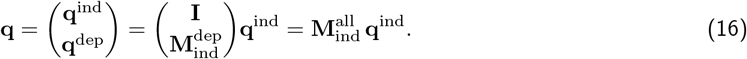

(ii) The thermodynamic forces are computed by the linear formula

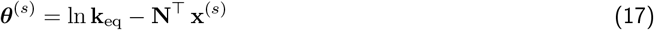

or briefly ***θ*** = **M**^*θ*^ **y** with a matrix **M**^*θ*^ obtained from the network structure. (iii) Based on rate laws and using Eq. (1), the enzyme concentration vectors **e**^(*s*)^ are given by

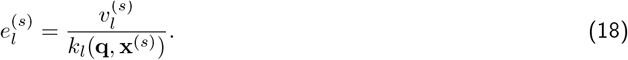

Both 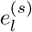 and its logarithm 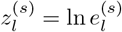 are convex functions in (**q, x**) for a wide range of rate laws. Moreover, the formula (18) yields positive enzyme concentrations on the entire solution space **x** (see below), except when fluxes vanish exactly 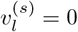. In this case, we directly set 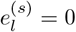 instead of estimating 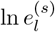 (where taking the logarithm would lead to problems.
3. **Solution space** The solution space for our free variables is defined by two sorts of constraints. First, lower and upper bounds on all variables except for the enzyme concentrations^27^

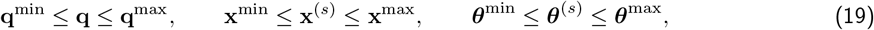

where *s* denotes metabolic states. Second, the driving forces must be positive (in the direction of the fluxes, and so the known flux directions define the signs of all driving forces. For all reactions with non-zero fluxes 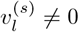, this yields the thermodynamic constraints

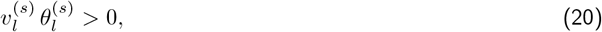

which (together with Eq. (17)) translate into linear constraints for the variable vector **y**. In reactions with zero fluxes, the driving forces are unconstrained (unless we know, for some other reasons, that reactions are in chemical equilibrium). Together all these constraints can be written as

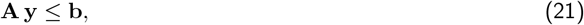

with a matrix **A** and a vector **b** obtained from reaction stoichiometries and flux directions. These constraints define a convex solution polytope *𝒫*. Each polytope point **y** describes a feasible set of model parameters and metabolic states (i.e. states with positive driving forces). Conversely, any feasible set of kinetic constants and metabolic states (respecting all bounds and constraints) corresponds to a point in the polytope.
4. **Priors and likelihood terms** The posterior is obtained from prior and likelihood terms. Kinetic constants, metabolite concentrations, and enzyme concentrations are considered in log-scale and described by Gaussian priors and likelihood terms on this scale. The log kinetic constants depend on a smaller set of independent kinetic constants. For these independent kinetic constants, we use a correlated prior arising from independent prior terms (for independent kinetic constants) and from pseudo values (for dependent kinetic constants). While pseudovalues are invoked formally to define correlated priors, they can be treated in practice as additional data points (see [15]). For the state variables (metabolite log-concentrations 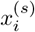 and enzyme concentrations 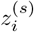), we assume uncorrelated Gaussian priors (i.e. log-normal priors for the concentrations themselves). Similarly, the logarithmic data values 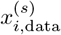 and 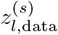 in the likelihood term, are assumed to be independent and normally distributed around the true values.

In contrast to Parameter Balancing and ECM, model balancing determines the vectors **q** and **x** at the same time. The resulting vector **y** lives in a high-dimensional polytope whose geometric structure is schematically shown in Figure 11. Since each state vector **y** consists of a vector **q** and a number of vectors **x**^*s*^, the polytope resembles a Cartesian product of the polytopes for these single vectors. However, thermodynamic constraints between kinetic constants and metabolite concentrations require that some parts of this Cartesian product must be removed. To see how the metabolite spaces for several metabolic states are combined, let us return to our simplified model balancing problem from section 2.3. We can solve this problem separately for each state, and this is also the easiest way to solve the problem. But we can also fit all metabolic states simultaneously by one big regression model. Each metabolite profile **x**^(*s*)^ must lie in a metabolite polytope 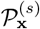. If the flux directions in all metabolic states are the same, these polytopes are identical; if fluxes change their directions, the metabolite polytopes 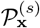 will differ. If we merge all vectors **x**^(*s*)^ into a vector **x**, the solution polytope for this vector will be higher-dimensional and will be given by the Cartesian product 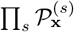. As before, we can consider the prior, likelihood, and posterior (for all metabolic states) as functions on this solution space. Since the metabolic states are independent, the prior, likelihood, and posterior functions can be split into products of priors, likelihoods, and posteriors for the single states, confirming that the estimation problems could have been solved separately.

### A.2 Convexity proof

To show that the truncated score function is convex, we note:

1. The logarithmic reaction time 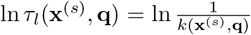 (for each reaction *l* and each metabolic state *s*) is convex in (**x**^(*s*)^, **q**) in the entire feasible space for (**x**^(*s*)^, **q**) and for all separable rate laws. For **x**^(*s*)^ alone, this has been shown in [39]. For the combined vector (**x**^(*s*)^, **q**), convexity is guaranteed by the fact that the formula for 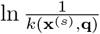, with separable rate laws, can be entirely decomposed into functions allowed in Disciplined Convex Programming.
2. The enzyme log concentration 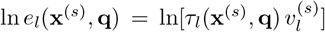 is linear in *τ*_*l*_(**x**^(*s*)^, **q**) and therefore a convex function in (**x**^(*s*)^, **q**).
3. In the prior score, likelihood score, and posterior score

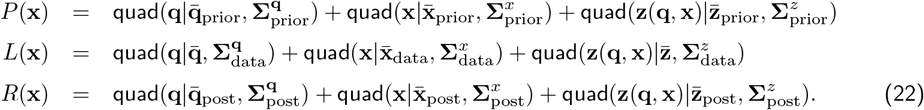

the first terms of are quadratic in **q** and independent of **x** and therefore convex in (**x**^(*s*)^, **q**), and the second terms of are quadratic in **x** and independent of **q** and therefore convex in (**x**^(*s*)^, **q**). The third terms, by themselves, are quadratic in **z**(**q, x**) and not necessarily convex. However, if they are are replaced by the truncated version, each of them can be written as a sum of non-decreasing functions of 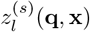, which are convex in (**q, x**), and are therefore convex in (**q, x**) themselves.

## B Implementation

A Matlab implementation of Model Balancing, together with example models and data, is available at https://github.com/liebermeister/model-balancing. The file format for models and data (kinetic constants, fluxes, metabolite concentrations, protein concentrations) is SBtab [48] and metabolic networks can be defined in SBML [49] or SBtab files. By default, the algorithm starts by running model balancing on an average model state (with metabolic state data given by the geometric mean over the metabolic states). The resulting kinetic constants are then used as initial values for the following full calculation with several metabolic states.

### B.1 Prior distributions

Since the values of non-identifiable parameters will be predicted based on their priors, the choice of realistic priors is important! Of course, the usage of priors affects all predicted variables, pushing them towards the prior means: high values will be estimated too low and low values will be estimated too high. This effect does not matter if a variable has been measured precisely or if priors are very broad. However, for non-identifiable variables the effect is large: while model balancing predicts a single (posterior mode) value, this value is almost arbitrary, determined entirely by the choice of the prior, and carries a large uncertainty (marginal posterior variance). On the one hand, this calls for posterior sampling (ninstead of reporting only the posterior mode). On the other hand, this underlines the need for realistic, informative priors. To define informative priors for kinetic constants [14, 50], we followed the procedure from parameter balancing [15] which takes into account parameter dependencies. We start by assuming independent Gaussian priors for all independent constants and (independent Gaussian) pseudo values for dependent constants: taken together, they define a correlated prior for all constants. Non-diagonal covariance matrices for ln *K*_eq_ values, obtained from equilibrator, could be taken into account. Mean values and standard deviations were taken from [18] and manually refined to match empirical distribution of kinetic constants. For the state variables (metablite and enzyme log-concentrations), we typically use diagonal covariances 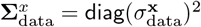 and 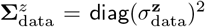, assuming independent measurement errors. However, correlated errors (e.g. caused by normalisation for metabolomics or proteomics samples) may be taken into account. There are several possible reasons to do this: first, concentrations differ more strongly between metabolites than between metabolic states for the same metabolite. This means, for the vector of all metabolite data, that values for each metabolite tend to be correlated. We could describe this by splitting the data values into 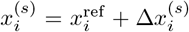, with a reference value 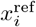 and deviations 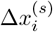. Assuming uncorrelated priors for 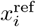 and 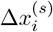 entails, directly, a correlated prior for 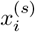 (and the same for enzyme log concentrations *z*_*l*_). Second, if we model metabolic time series, and assume that state variables vary smoothly over time, we can implement this assumption by imposing a prior that assumes correlations between (one and the same concentration at) subsequent time points. Third, there may even be reasons to assume correlated errors in data values: for example, in metabolomics data, differences in sample preparation or a normalisation per sample will lead to correlated measurement errors (by which, for example, all metabolite concentrations in one sample may appear too high or too low). In theory, such effects could be accounted for by assuming a correlated covariance matrix in the likelihood terms.

To define priors in practice, pseudo values, and constraints (for kinetic constants, metabolite concentrations and enzyme concentrations), we used the default values from parameter balancing (see www.parameterbalancing.net. However, when running parameter balancing as a test, we found that the available *k*_cat_ values were typically much higher than the prior median value as expected for enzymes in central metabolism [12]. In view of these data, we changed the prior for *k*_cat_ values from a median of 10 s^*−*1^ (geometric standard deviation 100) to a median of 200 s^*−*1^ (geometric standard deviation 50). Likewise, we changed the prior width for *K*_M_ values from a geometric standard deviation of 10 to a geometric standard deviation of 20 (while keeping the median 0.1 mM unchanged). A table describing the priors is provided in the github repository, file resources/data/data-prior/model-balancing prior.tsv. These values, used in the matlab implementation, can easily be modified. Importantly, in the actual model balancing calculations, a much broader prior was used in order to put more weight on existing data.

### B.2 Model balancing variants and simplifications

Model balancing can be adapted in various ways. (i) If a type of data is not used, likelihood terms for this data type are omitted. Even without any data, priors will keep the results in biochemically plausible ranges. (ii) If certain parameters (e.g. the equilibrium constants) are precisely known, their values can be predefined (e.g. by treating them as data with very small standard errors). (iii) Model balancing also applies to models with irreversible rate laws. In an irreversible rate law, there are fewer kinetic constants (since backward catalytic constants, equilibrium constants, and velocity constants do not play a role); the forward kinetic constant is a free parameter, and no Haldane relationship is considered. Describing (some or all) rate laws as irreversible changes the structure of the kinetic dependency matrix **M**. (iv) Different model parameterisations: instead of independent equilibrium constants, standard chemical potentials may be used as independent parameters [15]. (v) A preposterior for kinetic parameters may be obtained by previous parameter balancing, and pseudo values for metabolite and enzyme concentrations may be obtained by a previous ECM. (vi) To penalise unrealistically high metabolite or enzyme concentrations, a regularisation term may be added, for example, proportional to the cost function considered in ECM. (vii) Omics data may not contain absolute metabolite and enzyme concentrations, but relative changes between metabolic states. To account for such data, a variant of the dependency schema might be considered: for each metabolite, we split the log-concentrations 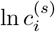 into a reference value ln *c*_*i*_ and a deviation 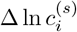. Uncorrelated priors for these variables yield a meaningful correlated prior for the metabolite concentrations, and a similar splitting can be used for enzyme concentrations.

So far we assumed that one data value is available for each of the model variables. For example, we assume that each metabolite level in each metabolic state has been measured, and that an *in-vitro* value is available for each kinetic constant. To account for the fact that values may be missing or measured multiple times, we need to relate a vector of model variables to the corresponding vector of variables measured (possibly, comprising multiple measurements of the same variable). The connection can be made by a data mapping matrix **D**_x_ (for metabolite log concentrations, and similar mapping matrices for other variables) that maps the vector of metabolite log-concentrations in the model onto the vector of metabolite log-concentration data values. In the ideal case (all values have been measured exactly once), **D**_x_ is a simple identity matrix. If a concentration has not been measured, the corresponding row of this identity matrix must be omitted, and if a value has been measured multiple times, the row must be duplicated. With mapping matrices, our likelihood loss score (with terms for kinetic constants and state variables) becomes

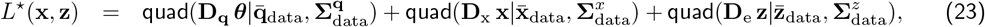

and formula (7) for posterior means and covariances reads

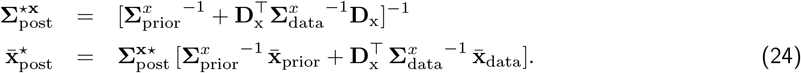

Of course, missing and multiple values for kinetic constants, enzyme concentrations, and thermodynamic forces can be treated in the very same way (with analogous terms for **q, z**, and ***θ***). This is particularly important for kinetic constants, where typically only a small fraction of the constants have been measured *in vitro*. Finally, instead of using mapping matrices, there is also a simpler practical solution. Whenever data values are missing, we can replace them by invented values with infinite error bars. Likewise, if multiple data values exist for one variable, we can merge them into a single value to be used as a data point. The calculation follows the same scheme a Eq. (24), with a vector 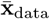 containing the multiple measured data values, *σ*_*xl*_ their standard deviations, **D** = (1, 1,..)^*T*^ (relating a single model variable to multiple measured values), and no prior. We obtain

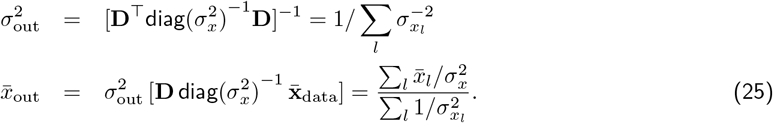

### B.3 Practical computation details

1. **Calculation of the posterior** To compute the posterior score from prior and likehood scores (Eq. 7 for metabolite concentrations, and similar formulae for enzyme concentrations and kinetic constants), we need to invert a covariance matrix. This can be numerically expensive. To compute the posterior of the independent kinetic constants, we need to solve

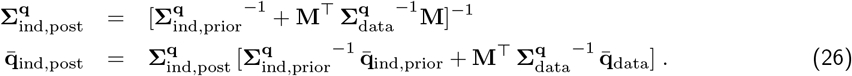

The calculation of 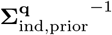 and 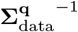 (precision matrices for metabolite and enzyme concentrations) is easy because the original covariance matrices are diagonal and the mapping matrices **D** select single vector elements. However, inverting the term in brackets may be hard. To speed up the calculation, we set 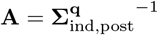 and obtain the similar formulation

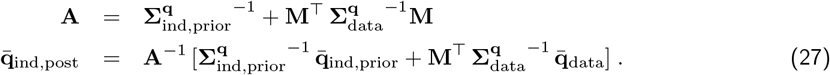

Now the costly matrix inversion in the first equation is avoided, and the right-hand side in the second equation can be computed without explicitly computing the matrix inverse (e.g. by using the matrix left division operator \ in matlab). This calculation is faster and works for sparse matrices.
2. **Reactions with vanishing flux** If reaction flux is non-zero, the flux direction puts a constraint on the driving force, and the predicted enzyme concentration is positive. If a reaction is always inactive – that is, in *all* metabolic states – the kinetic constants for this reaction are ill-determined, and the reaction can be removed from the model. But what if a reaction fluxes vanish in some of the metabolic states? The vanishing flux can either be caused by a vanishing enzyme concentration, or by a vanishing thermodynamic force. If the reaction is known to be in chemical equilibrium, we also set the driving force to 0, which leads to an extra equality constraint on metabolite concentrations. In this case, the enzyme concentration can be positive and needs to be estimated (although the economical “principle of dispensable enzyme” would suggest a vanishing enzyme concentrations in this case). Otherwise, with a zero flux and a non-zero driving force, the enzyme activity must be zero: for an enzyme without allosteric inhibition, this means that the enzyme concentration must vanish.
3. **Divergence of enzyme concentrations close to polytope boundaries** Each thermodynamic constraint defines a boundary of the solution polytope. Close to this boundary, an enzyme concentrations goes to infinity and the likelihood function explodes. This steep increase can cause numerical problems during optimisation. To handle them, we may apply the logarithm function once more to the (likelihood or posterior) score, and use the resulting function as our minimisation objective. This new objective function will still go to infinity at polytope boundaries, but less steeply. The new objective function may be non-convex, but since it depends monotonically on a convex function, it will still have a single local minimum. A second way to avoid this problem is to exclude problematic regions close to the boundary by introducing some extra constraints. In practice. we can make all thermodynamic constraints a bit tighter, by requiring small, non-zero thermodynamic forces in every reaction [39].
4. **Starting point for optimisation** To obtain an initial point for our optimisation, we may first run model balancing for an average metabolic state. This yields a first guess of the kinetic constants. Alternatively, we can run model balancing separately for each metabolic state. In each run, we start from the prior mode (or alternatively, from the posterior mode for kinetic constants obtained by Parameter Balancing, and the posterior mode for each metabolite value). The resulting concentration vectors and the state-averaged (arithmic/geometric) kinetic constant vector can be used as initial values for the multi-state problem.
5. **Running parameter balancing as a separate first step** Model balancing can also be run in two steps. The first step, is a simple parameter balancing problem: we consider only kinetic constants and fit them to kinetic data. The result is a multivariate Gaussian posterior for all log kinetic constants [15] that summarizes all data and prior knowledge about the kinetic constants. In the second step, we use this posterior as a prior for the kinetic constants, and fit kinetic constants and model states (metabolite and enzyme concentrations) to metabolite and enzyme data. Since the kinetic data have already been used to define the prior, they can be ignored in this part of the estimation. The calculation is equivalent to the method described in this paper. By processing the kinetic data separately in advance, we can learn more clearly what information is contained in the kinetic data alone, before combining them with metabolic data. Moreover, a known kinetic “prior” that includes all information about kinetic data may allow us to further constrain the kinetic constants in order to reduce the solution space.

## C Example model

### C.1 Model structure

The *E. coli* central carbon metabolism model from [39] comprises 40 metabolites and 30 reactions and contains 107 *K*_M_ values and 167 kinetic constants in total (*K*_M_ values, as well as forward and backward *k*_cat_ values). The model structure is shown in Figure 12 and described at https://github.com/liebermeister/model-balancing (in the file resources/data/data-organisms/escherichia coli/network/ecoli noor 2016.tsv).

To model aerobic growth on glucose, we used state data from [39] which gather measured flux data from [51], proteomics data from [52], and metabolomics data from [53]. To model several metabolic states, we used a data set from [20], where a larger network model had been considered, proteomics data from different sources were used, and flux data had been computed by FBA. I linearly the flux data onto the *E. coli* model to obtain complete and consistent flux values. A comparison between the two data sets reveals a discrepancy in scaling: the (FBA-derived) fluxes from [20] were smaller than the fluxes taken from [39] by an approximate factor of 10, while enzyme concentrations were smaller by an approximate factor of 2.

### C.2 Artificial data

Artificial kinetic constant data were generated as follows. Given the network structure, true artificial kinetic constants were generated by assigning random (log-normal) values to 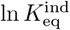, ln *K*_*M*_, and ln *K*_*V*_ and computing the other constants. The random values were sampled from the same distributions that are used as priors in model balancing.

To generate artificial metabolic state data, enzyme concentrations and external metabolite concentrations were randomly sampled from the same distributions that are used as priors in model balancing. Then the model was parameterised with the “true” artificial kinetic constants and was solved to obtain a steady reference state (steady-state metabolite concentrations and fluxes). Based on this reference state, a number of metabolic states were constructed by randomly varying metabolite and enzyme concentrations (again, following the prior distribution) and computing the (non-steady) reaction rates^28^. The resulting states are seen as the “true values”. To generate noisy state data, uncorrelated random noise was added to the “true values”. When generating artificial data, noise was also added to fluxes but the flux signs were kept unchanged, to ensure thermodynamically feasible flux directions as required in model balancing.

## Footnotes

1 Note that kinetic models typically use rate laws such as Michaelis-Menten kinetics that are based on quasi-steady state approximation on an even faster time scale.

2 Equilibrium constants are determined by thermodynamics and do not depend on specific enzymes, but for simplicity we will also call them kinetic constants.

3 For ease of terminology, kinetic constants and state variables will be called “model variables” (even though kinetic constants would usually be called “parameters”, and the methods for estimating our model variables are usually called “parameter fitting”).

4 Mathematically, this estimation problem resembles Enzyme Cost Minimisation (ECM) [39]. Both methods are based on kinetic models with known parameters and predefined fluxes, and both of them optimise metabolite and enzyme concentrations, but in different ways. In ECM, metabolite and enzyme concentrations are optimised for a minimal biological cost, while in the present estimation problem, they are fitted to data.

^5^

6 Exact zero concentrations are deemed impossible, except for enzyme concentrations in the case of exactly vanishing fluxes, which are not estimated but directly set to zero).

7 Typically, we assume uncorrelated priors, with covariance matrices 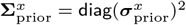.and 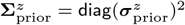

8 Negative metabolite concentrations are automatically excluded by optimising metabolite concentrations in log scale. For enzymes, negative concentrations are excluded by restricting the metabolite concentrations to a thermodynamically feasible solution space, where the enzyme demand formula is guaranteed to yield positive results.

9 A formula for the general case with missing and duplicate data values is given in appendix B.2.

10 Note that the precision matrices (inverse covariance matrices) are additive, and that the posterior mean is a weighted sum of prior and likelihood means, with “matrix prefactors” 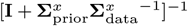 and 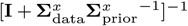

11 A term for thermodynamic driving forces would also be convex because it is composed of a convex function quad() and a linear function ***θ***(**x**), which together yields a convex function.

12 A similar situation exists in ECM, where upper bounds on (absolute) enzyme and metabolite concentrations (or their weighted sums) are convex constraints in **x** and can be considered in the optimality problem, while lower bounds (as well as equality constraints) on the same quantities are non-convex constraints and cannot be used.

13 The preposterior for kinetic constants is given by the posterior obtained from parameter balancing. For more details, see appendix B.3.

14 Pseudo values are a way to define priors by which all model parameters, even dependent ones, have non-flat priors. Details can be found in appendix A.2.

15 The bottom subfigure rows (estimation without kinetic data) are repeated between Figures 3 and 4, and accordingly between other figures.

16 The geometric standard deviation is defined as exp(*σ*), where *σ* is the root mean square of the residuals on (natural) log-scale.

17 In theory, the predictive capacity of model balancing for kinetic constants could be studied by a leave-one-out crossvalidation (where all in-vitro kinetic constants except one are used as data, and the remaining value is used as validation data. However, it is to be expected that the results will lie in between the results of the “fitting” (top row) and “prediction” scenario (center or bottom row).

18 In the SIMMER method [21], a Markov chain Monte Carlo approach is used for the optimisation. The estimation can be reformulated as a model balancing problem.

19 Measurement errors in metabolic fluxes will distort our estimation results, but model balancing remains applicable. However, fluxes must be thermodynamically consistent, that is, without thermodynamically infeasible flux cycles.

20 Accordingly, kinetic constants and metabolite concentration must be described with log-normal distributions for measurement errors and priors while enzyme concentrations must be described on non-logarithmic scale (assuming normal distributions for measurement errors and priors).

21 Mathematically, parameter balancing resembles the component contribution method, which component contribution method [46] used to determine thermodynamic constants in eQuilibrator [47].

22 The equilibrium constants were not parameterised by standard chemical potentials *µ*^*°*^ (as proposed in [18] for parameter balancing), but by independent equilibrium constants. This is convenient because we use a smaller set of independent variables and avoid non-identifiability (while the standard chemical potentials themselves are not in the centre of interest), and the same choice could be applied in parameter balancing.

23 To make use of rate laws and flux data in parameter balancing, a trick can be used: the balanced kinetic constants can be adjusted to match the fluxes. However, this only works for a single metabolic state, and information from several states cannot be aggregated.

24 Many metabolic modelling methods, such as FBA, assume stationary flux distributions: model balancing does not make this assumption. Like ECM it applies to non-stationary fluxes, e.g. fluxes appearing in dynamic time courses.

25 In the matlab implementation, this is automatically checked.

26 In parameter balancing, the independent parameters behind the equilibrium constants are Gibbs free energies of formation. The Gibbs free energies of formation determine all equilibrium constants, but in a “redundant” way. While they could be used in model balancing too, an alternative (minimal) set of variables is given by the equilibrium constants of a subset of the reactions *l*, where the columns of the stoichiometric matrix (for all external and internal metabolites) corresponding to these reactions form a maximal set of linearly independent columns.

27 For enzyme concentrations, positivity is ensured by the other formulae. Upper bounds on individual enzyme concentrations (or on weighted sums of enzyme concentrations) would define non-linear inequality constraints that remove some regions from the solution polytope, giving it a curved shape. However, since these constraints are convex, the resulting solution set would still be convex. Such constraints are not considered here, but could be added without any problems. Lower bounds on enzyme concentrations (or on absolute metabolite concentrations), in contrast, would make the solution space non-convex.

28 Alteratively, one could simulate a dynamic time course and take snapshots (fluxes, metabolite concentrations and enzyme concentrations) at different time points.

